# The compositional nature of the human alpha rhythm

**DOI:** 10.64898/2026.01.24.701499

**Authors:** Maëlan Q. Menétrey, David Pascucci

## Abstract

Resting-state brain activity is dominated by oscillations in the alpha band (8–13 Hz). The dominant frequency within this band, the individual alpha peak frequency (IAPF), has been widely used as a global marker of inter-individual variability in perceptual and cognitive functions, as well as clinical phenotypes. However, the traditional approach implicitly assumes a single oscillatory rhythm varying along a parametric continuum. Here, we analyzed resting-state EEG data from more than 2000 participants across multiple independent datasets using unsupervised spectral decomposition methods. We identify three distinct and highly stable alpha components, or archetypes, that jointly and compositionally determine the IAPF. These components exhibit dissociable age-related trajectories as well as distinct scalp topographies and cortical sources. Together, these findings indicate that individual differences in alpha activity do not reflect variation along a single frequency dimension, but rather differences in the relative contribution of discrete alpha generators. These results call for a reevaluation of many reported associations between alpha frequency, brain function, and clinical phenotypes, and suggest that estimates of alpha-component expression may provide more precise and informative biomarkers than conventional global IAPF measures.

## Introduction

Brain activity is dominated by oscillations in the 8–13 Hz range, commonly referred to as alpha activity (Figure 1A) ^[1]^. Over the past century, parameters of these oscillations, and in particular the individual alpha peak frequency (IAPF, i.e., the dominant frequency within the alpha band ^[2]^), have emerged as candidate biomarkers for predicting brain function and dysfunction ^[3]^.

In non-invasive scalp electroencephalography (EEG), IAPF is typically estimated as an average alpha peak across all electrodes or a subset (Figure 1B). This measure exhibits a striking degree of inter-individual variability ^[4]^ alongside strong test–retest reliability ^[5,6]^, making it a potential neurophysiological trait-like marker with both theoretical and clinical value. Indeed, variability in IAPF has been linked to inter-individual differences in a variety of perceptual ^[7–10]^ and cognitive tasks ^[11–14]^, as well as to general cognitive traits ^[15]^ and clinical conditions ^[16,17]^. In particular, a systematic slowing of IAPF has been reported in conditions such as cognitive decline ^[18,19]^, epilepsy ^[20]^, autism ^[21]^, Alzheimer’s and Lewy body dementia ^[22,23]^, Parkinson’s disease ^[24]^, schizophrenia ^[8,11,25]^ and depression ^[26]^.

The heterogeneity of reported associations and the limited understanding of the neural mechanisms underlying scalp IAPF, however, have rendered it a largely transdiagnostic marker, with limited functional and diagnostic specificity and weak causal insight ^[3]^. For example, it remains unclear whether inter-individual variability in IAPF reflects parametric variation within a single dominant generator (Figure 1C) or instead arises from the superposition of multiple alpha generators with distinct anatomical and functional roles, as suggested by several lines of research ^[27–35]^. If multiple alpha generators are involved, then the single scalar estimate of global IAPF, as traditionally used, provides only a coarse and ambiguous summary, conflating many different combinations of underlying alpha components into the same value. Such ambiguity may contribute to the limited specificity of global IAPF measures and the heterogeneity of reported findings, and calls for a reassessment of how alpha activity, and in particular frequency-related markers, are quantified, interpreted, and linked to behavior and pathology.

Here, in a large-scale sample of task-free EEG recordings drawn from multiple resting-state datasets totaling over 2000 participants, we tested the hypothesis that alpha activity, and therefore the estimated IAFP, is composed of multiple, partially independent components. Using unsupervised spectral decomposition, we assessed whether distinct alpha components with specific scalp topographies and neural generators coexist within individuals, and evaluated their contribution to inter-individual variability in IAPF and age ^[36–38]^, as well as their longitudinal stability, and their cortical sources. Our results reveal highly stable and reproducible alpha “archetypes”, i.e., components that are systematically conflated in conventional global IAPF estimates. Each individual expresses a characteristic mixture of these alpha components, defining a composite neural trait analogous to a fingerprint. We argue that variation in the relative weighting of these components can substantially alter the interpretation of IAPF, motivating a new framework for alpha “profiling” that enables more mechanistic and reliable links between alpha rhythms, behavior, brain function, and clinical outcomes. Within this framework, we further propose how the method can be used to derive single-participant estimates of component expression, which in this study show predictive value for age and may therefore constitute a promising biomarker approach beyond IAPF alone.

## Results

### Resting-state EEG is characterized by a mixture of archetypical alpha components

We estimated the average scalp power spectral density (PSD) from resting-state EEG data in a sample of 1268 participants spanning the age range from 5 to 83 years (TDBRAIN dataset ^[39]^). The PSD exhibited the typical pattern observed in resting-state EEG, characterized by a dominant peak in the alpha band and a gradual decrease in power with increasing frequency, consistent with the so-called aperiodic (1/f-like ^[40]^) component (Figure 1A). When IAPF was estimated at the population level using a conventional global scalp approach, thus, implicitly assuming a single dominant generator, the mean IAPF across participants was 9.62 Hz (SD = 1.07), with substantial inter-individual variability (Figures 1B-C).

To test whether global IAPF instead reflects a combination of multiple alpha components, we applied two complementary unsupervised spectral decomposition approaches. First, we performed a frequency-domain principal component analysis (fPCA) ^[41–43]^, retaining components that explained at least 5% of the population-level variance (see Methods). This analysis revealed three distinct components (i.e., factor loadings, corresponding to frequency profiles) within the alpha range, corresponding to *slow* (fPC2), *middle* (fPC3), and *fast* (fPC4) alpha components, alongside a low-frequency, non-alpha component (fPC1; Figure 1D). Together, these components accounted for 80.6% of the variance (26.6% for fPC1, 16.7% for fPC2, 21.4% for fPC3, and 16.0% for fPC4). In simulations, we assessed the performance of fPCA under different scenarios (Supplementary Material “Simulations” and Figure S1) and used an additional complementary approach to assess whether the identified components reflect distinct spectral patterns within individuals (Supplementary Material “Non-negative matrix factorization” and Figure S2).

**Figure 1.**
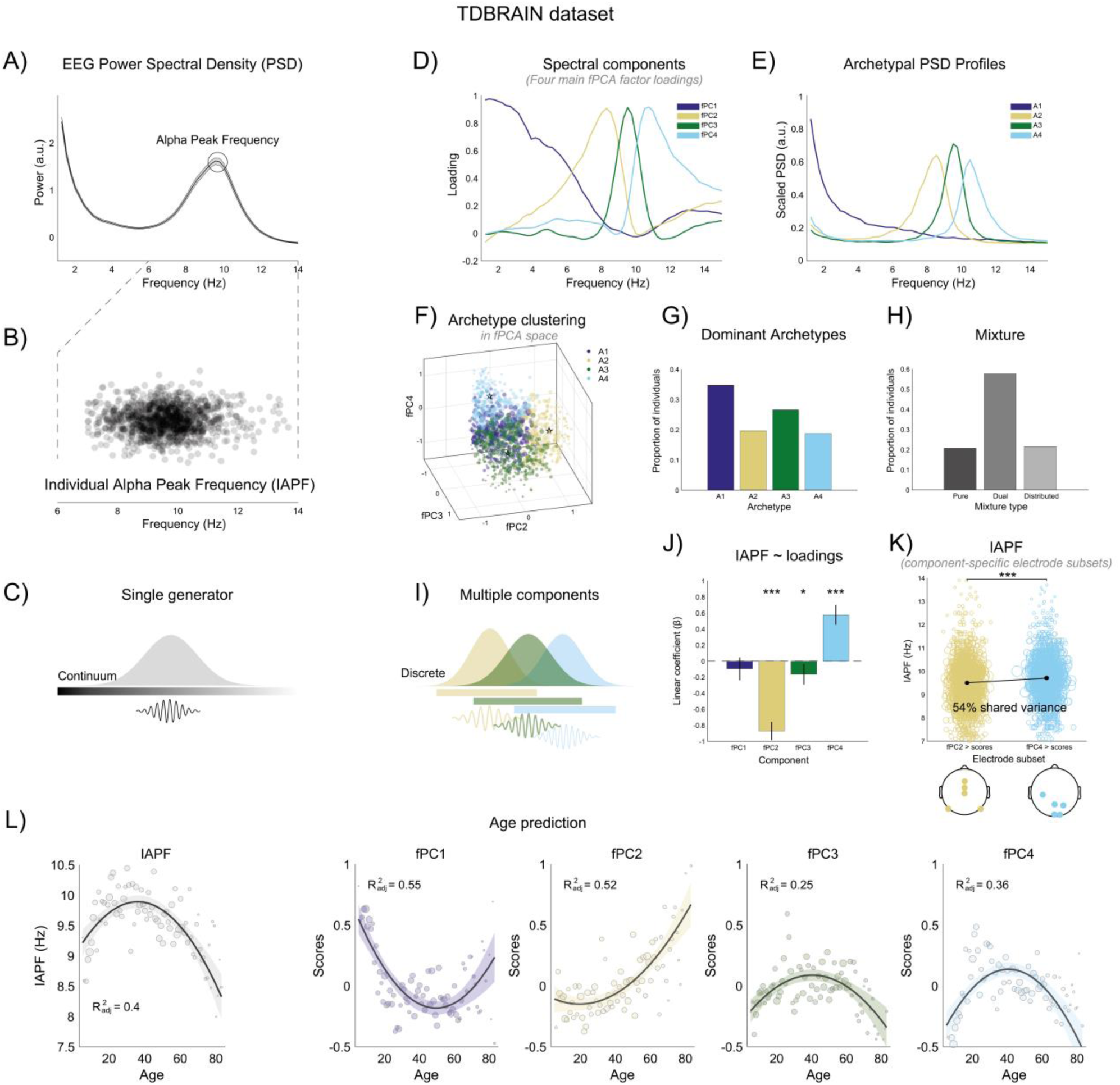
Resting-state EEG is characterized by a mixture of archetypical alpha components. A) EEG power spectral density (PSD) typically exhibits an alpha peak (increased power around 10 Hz), commonly summarized by the individual alpha peak frequency (IAPF). B) IAPF varies widely across individuals (each dot represents one participant from the TDBRAIN dataset ^[39]^). C) An overly simplified account depicting IAPF as a single, unified index reflecting the activity of a single generator varying along a continuum of oscillatory frequency, or “speed”. D) Frequency-domain principal component analysis (fPCA) identified four main spectral components explaining ≥80% of the variance, including three components within the alpha band. E) Archetypal analysis identified four archetypes explaining ≥95% of the variance, with spectral profiles closely matching those recovered by fPCA. In panels D and E, the slow (fPC2), middle (fPC3), and fast (fPC4) alpha components are shown in sand, dark green, and light cyan, respectively, whereas the slower non-alpha component is shown in dark purple. Both decompositions were performed on the TDBRAIN dataset (N = 1268). F) Three-dimensional fPCA score plot (fPC2–fPC4), with individual participants projected in fPCA space. Component scores (also termed factor scores in PCA) quantify the expression of each component and are obtained by projecting electrode-wise spectral profiles onto the fPCA loading patterns, as in standard PCA. To obtain a stable single-participant estimate, we computed an aggregate score magnitude by averaging electrode-wise scores within each participant, thereby summarizing the overall expression of each component across the scalp. Participants are then color-coded according to their dominant archetype, from archetypal analysis. Dot size is scaled by the participation ratio (see Methods), with larger dots indicating a more distributed contribution of multiple components. Archetype centroids are indicated by stars. G) Proportion of participants for whom each archetype had the highest weight, indicating the dominant archetype in each individual’s spectral profile. H) Distribution of participants across the three participation-ratio categories: pure (dark gray), dual (medium gray), and distributed (light gray), reflecting increasingly distributed contributions across archetypes. I) The patterns identified by fPCA and archetypal analysis suggest that global IAPF estimates reflect a mixture of multiple alpha-related spectral components with distinct peak frequencies, rather than a single, unified measure of alpha activity. J) IAPF, estimated from the average scalp PSD, was significantly predicted by the individual scalp-averaged scores associated with the slowest and fastest alpha spectral components (i.e., Components 2 and 4 in Figure 1D), with effects in opposite directions, with a weaker contribution from the mid alpha component. In other words, IAPF was predicted by the relative expression of two alpha components across individuals. Error bars represent 95% confidence intervals for the linear coefficients (β) estimates. K) IAPF estimated from electrode subsets showing the strongest scores for the slow (sand) and fast (light cyan) alpha components. The two estimates differed significantly (\*\*\**p* < .001), with 54% of shared variance. Dot size is inversely scaled to each participant’s distance from the population mean, with larger dots indicating values closer to the mean. The head plots below show the electrode subsets selected for each component. L) Relationship between age and global IAPF (first panel) and relationships between age and the four spectral components (last four panels; component scores averaged across electrodes for each participant). Age-related variance was associated with changes across all four components, suggesting that age-related changes in global IAPF reflect shifts in the relative expression of multiple spectral components rather than uniform slowing. The non-alpha and slow alpha components also provided a better model of age than IAPF alone, indicating that component-specific measures capture age-related information not fully reflected in the global IAPF estimate. The solid line and shaded area show predictions from a quadratic model with 95% confidence intervals.

As a complementary approach that does not impose orthogonality constraints, we applied archetypal analysis ^[44,45]^, an unsupervised method that represents each individual in a dataset as a mixture of “pure type” profiles, to identify a set of archetypal spectral patterns expressed in varying proportions across participants. This analysis also identified four archetypes that explained 95.7% of the variance. The resulting archetypal spectral profiles closely resembled those recovered via fPCA (Figure 1E), as indicated by strong correlations with the corresponding spectral loadings (slow alpha: *r* = .87; middle alpha: *r* = .80; fast alpha: *r* = .77; low-frequency non-alpha: *r* = .86). These profiles were expressed broadly across the population (Figure 1F), with a slightly higher proportion of individuals showing maximal expression of the low-frequency non-alpha component and the middle alpha component (Figure 1G).

We then quantified, for each participant, the effective number of archetypes contributing to their spectral profile defining a metric termed the participation ratio (see Methods). Participants were classified as showing predominantly single-archetype (pure), dual-archetype, or distributed profiles, reflecting increasing contributions from multiple archetypes. Most participants (>50%) showed dual-archetype profiles, while approximately 20% each showed predominantly single-archetype (pure) or distributed profiles (Figure 1H).

This pattern confirmed that individuals generally exhibit mixtures of these spectral profiles (Figure 1I), and that no single archetype dominates strongly for most participants. We further show participant subtypes to illustrate the different PSD patterns observed when at least two components contribute substantially to an individual’s spectral profile (see Supplementary Material “Within-individual electrode subtyping” and Figure S3).

### Individual differences in alpha components drive variability in IAPF and aging

The results from both decomposition methods converged, indicating that the global IAPF typically estimated at the scalp reflects a mixture of multiple alpha components rather than a single oscillator ^[27–35]^. Thus, global IAPF likely captures the relative contributions of these distinct components and their combination in each individual’s scalp EEG, rather than parametric variation of a unitary intrinsic rhythm.

To directly test this possibility, we modeled each participant’s IAPF (N = 1268; estimated with standard procedures from the average scalp PSD; see Methods) as a function of the scalp-averaged component scores obtained from our fPCA spectral decompositions (see Methods). IAPF was significantly predicted by the expression of slow and fast alpha components, with opposite regression weights (slow alpha component: *β* = -0.87, SE = 0.05, *t* = -14.91, *p* < .001; fast alpha component: *β* = 0.57, SE = 0.06, *t* = 9.15, *p* < .001), whereas the expression of the remaining components made smaller or no contributions (middle alpha component: *β* = -0.16, SE = 0.06, *t* = -2.49, *p* = .01; low-frequency non-alpha component: *β* = -0.09, SE = 0.07, *t* = -1.32, *p* = .18) (Figure 1J). On this basis, we constructed a simplified model using the difference between the slow and fast alpha scores to predict IAPF. This contrast significantly predicted IAPF (*β* = -0.73, SE = 0.02, *t* = -31.64, *p* < .001) and accounted for up to 44.1% of its variance. Thus, variability in IAPF is largely due to the relative expression of two distinct alpha components. Importantly, when IAPF was estimated separately from two subsets of five electrodes preferentially expressing the slow and fast alpha components, respectively (see Methods), the resulting estimates differed significantly (paired t-test, *t* = -9.35, *p* < .001, Figure 1K) and shared only 54% of their variance (R^2^ from a linear model). That is, IAPF estimates from the two scalp regions capture partly non-overlapping information that may be lost when a single global IAPF value is used.

To further prove this point, we focused on the known relationship between IAPF and age ^[46,39,38]^. As expected, age was a strong predictor of IAPF, and model comparison showed that a quadratic model provided a substantially better fit than a linear model (delta BIC = 34.3). In the quadratic model, both the linear age term (β = 0.05, SE = 0.008, *t* = 6.19, *p* < .001) and the quadratic term (β = −0.0007, SE = 0.0001, *t* = −6.93, *p* < .001) were significant, indicating a nonlinear trajectory in which IAPF increases during early adulthood, plateaus, and declines in older age (Figure 1L). This result, if taken blindly, would be suggestive of a general slowing of alpha rhythms with age ^[30,47,48]^.

However, when modeling age as a function of the expression (scalp-averaged component scores) of the four spectral components derived above, a more nuanced scenario emerged (Figure 1L). For both the fast and middle alpha components, quadratic models provided a better fit than linear models (fast alpha: delta BIC = 32.4; middle alpha: delta BIC = 19.9). In both cases, age showed significant positive linear effects and significant negative quadratic effects, consistent with inverted U-shaped trajectories (fast alpha, linear: β = 0.03, SE = 0.004, *t* = 6.79, *p* < .001; quadratic: β = −0.0003, SE = 0.00005, *t* = −6.71, *p* < .001; middle alpha, linear β = 0.02, SE = 0.003, *t* = 5.24, *p* < .001; quadratic: β = −0.0002, SE = 0.00004, *t* = −5.22, *p* < .001), indicating increases from early adulthood to midlife followed by declines at older ages. In contrast, the expression of the slow alpha component was best described by a nonlinear model showing an increasingly positive association with age (delta BIC = 11.1). While the linear term alone showed negligible effects (β = −0.007, SE = 0.004, *t* = −1.96, *p* = .08), the quadratic term was positive and significant (β = 0.0002, SE = 0.00004, *t* = 4.06, *p* < .001), consistent with a progressive increase later in life. The expression of the low-frequency non-alpha component also showed a nonlinear association with age (delta BIC = 31.5; linear term: β = −0.03, SE = 0.004, *t* = −8.13, *p* < .001; quadratic term: β = 0.0003, SE = 0.00005, *t* = 6.61, *p* < .001). Importantly, the expression of the slow alpha and low-frequency non-alpha components provided a better fit to age than IAPF alone (slow alpha: R^2^ = 0.52; low-frequency non-alpha: R^2^ = 0.55; IAPF: R^2^ = 0.40), demonstrating that component expression captures age-related variance not reflected in global IAPF.

Collectively, these results show that aging is not associated with a uniform slowing of global alpha activity, but rather with complex, component-specific changes in spectral contributions, including a growing dominance of the slow alpha component.

### Alpha components are stable individual traits, replicable across datasets

We ran the same analyses on an independent resting-state EEG dataset, the DVS dataset (N = 595; age range 20–70) from a longitudinal study ^[49]^, which includes repeated recordings on the same day and again after five years for a subset of participants. Using the same criteria and procedures applied to the TDBRAIN dataset, we recovered the four main spectral components from the fPCA factor loadings, which together explained 80.7% of the variance (27.5% for fPC1, 16.4% for fPC2, 22.4% for fPC3, and 14.4% for fPC4). The frequency profiles closely overlapped with those reported in the TDBRAIN dataset (Figure 2A, see also Figure S3), with comparable archetypal expression patterns (Figure 2B; see also Figure S4), NMF components (Figure S2), and similar relationships between individual-level expression of component scores, IAPF and age (Figures S5 and S6). As in TDBRAIN, the component score topographies identified distinct electrode subsets preferentially expressing the slow and fast alpha components. IAPF estimates from these subsets differed significantly (paired *t*-test, *t* = −8.45, *p* < .001) and shared only 43% of their variance (Figure 2C).

These results demonstrate that the spectral components are robust and replicable across independent datasets, recorded in a different laboratory, with different participants, hardware, and preprocessing pipelines.

**Figure 2.**
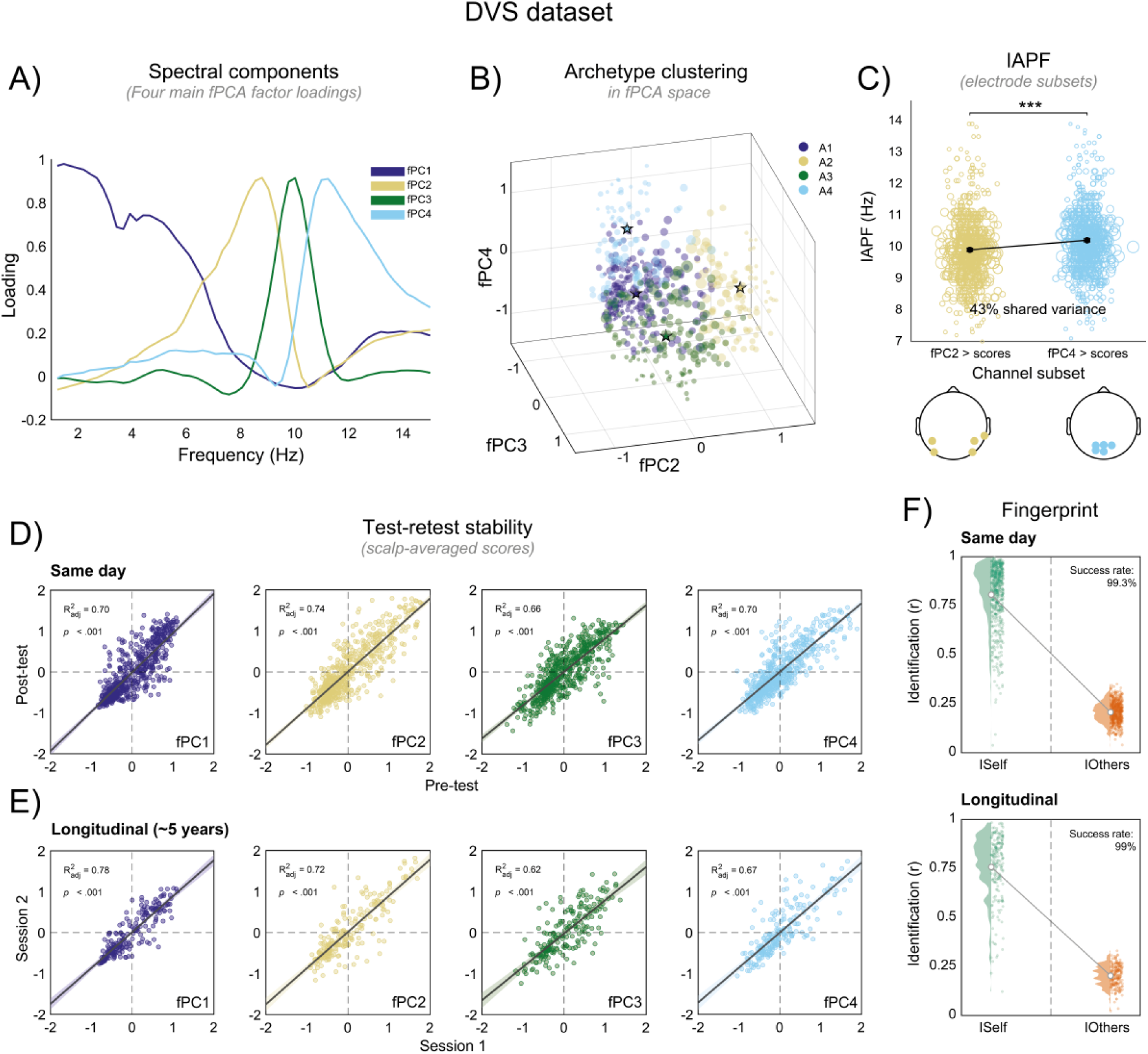
Alpha components are stable individual traits, replicable across datasets. A) In the DVS dataset (N = 595), fPCA identified four main spectral components explaining ≥80% of the variance, including three components within the alpha band. B) Three-dimensional fPCA score plot (fPC2–fPC4) as in Figure 1F. C) IAPF estimated from electrode subsets as in Figure 1K, showing the strongest scores for the slow (sand) and fast (light cyan) alpha components. The two estimates differed significantly (\*\*\**p* < .001), with 43% of shared variance. D) Short-term test–retest stability (N = 590), quantified as the correlation between scalp-averaged scores (i.e., scores associated with each fPCA loading, averaged across electrodes), estimated from EEG recordings separated by approximately 2 hours. The solid line and shaded area show predictions from a linear model with 95% confidence intervals. E) Long-term test–retest stability assessed over a 5-year interval (N = 202). F) Fingerprint analysis. Identifiability was quantified using within-subject correlations between component score topographies across sessions, both on the same day (upper plot) and five years apart (lower plot). Higher within-subject similarity (ISelf) relative to between-subject similarity (IOthers) indicates strong subject-specific reliability, demonstrating that the expression of the spectral components constitutes a stable and distinct individual fingerprint.

We then exploited the longitudinal structure of the dataset to assess the test–retest reliability of the expression of each spectral component (component scores). Applying the same spectral decomposition to each session, we quantified the stability of individual scalp-averaged component scores, that is, each component’s contribution to a participant’s scalp PSD (see Methods). The expression of all components showed strong reliability, both within the same day (two sessions separated by approximately two hours; Figure 2D; low-frequency non-alpha component: β = 0.96, SE = 0.02, *t* = 36.77, *p* < .001; slow alpha: β = 0.89, SE = 0.02, *t* = 40.98, *p* < .001; mid alpha: β = 0.81, SE = 0.02, *t* = 33.7, *p* < .001, fast alpha: β = 0.83, SE = 0.02, *t* = 36.69, *p* < .001) and across five years (for 202 participants; Figure 2E; low-frequency non-alpha component: β = 0.87, SE = 0.03, *t* = 26.44, *p* < .001; slow alpha: β = 0.88, SE = 0.03, *t* = 22.51, *p* < .001; mid alpha: β = 0.81, SE = 0.04, *t* = 18.25, *p* < .001, fast alpha: β = 0.85, SE = 0.04, *t* = 20.36, *p* < .001). All other results found in the TDBRAIN dataset and in the first session of the DVS dataset were also reproduced in the second session of the DVS dataset (Figures S2-S6).

At the individual level, the spatial pattern of component scores across electrodes was highly specific. A fingerprinting analysis (see Methods) comparing within-subject correlations across sessions with between-subject correlations revealed near-perfect identifiability (success rate = 99.3% in the same day, 99% after five years; Figure 2F). In other words, the topographies of component scores are highly stable over time and exhibit strong trait-like specificity at the individual level.

### Alpha spectral components map onto distinct spatial and cortical patterns

We replicated the results in another independent dataset, the LEMON dataset (N = 198; young: 20– 35; older: 59–77 ^[50]^), identifying the same four spectral components (Figure 3A; see also Figures S2 and S4), which together explained 83.1% of the variance (17.2% for fPC1, 18% for fPC2, 24.1% for fPC3, and 23.8% for fPC4). The expression of these components showed associations with IAPF and age comparable to those observed in the TDBRAIN dataset (Figures S5 and S6).

For the LEMON dataset, we here report the component score topographies, as a key test of the interpretation of distinct components (see Methods and Figure 3C). These maps show the fPCA scores associated with each component at each electrode, i.e., their expression at the scalp topography level, averaged across participants. The scalp distributions of the fPCA scores differed across components (Figure 3C). In particular, the slow alpha component was expressed more strongly over occipito-temporal electrodes, whereas the fast alpha component showed maximal expression over occipito-parietal electrodes (Figure 3D). We confirmed these distinct spatial patterns using a “back-propagation” regression approach that avoids any ambiguity in the interpretation of PCA scores and their sign (see Methods and Figure 3C). Comparable spatial patterns were also observed in all the other datasets (Figure S7; see also Figure S8 and Supplementary Material: “Replication of the four spectral components in a small sample dataset” for an additional resting-state EEG dataset).

**Figure 3.**
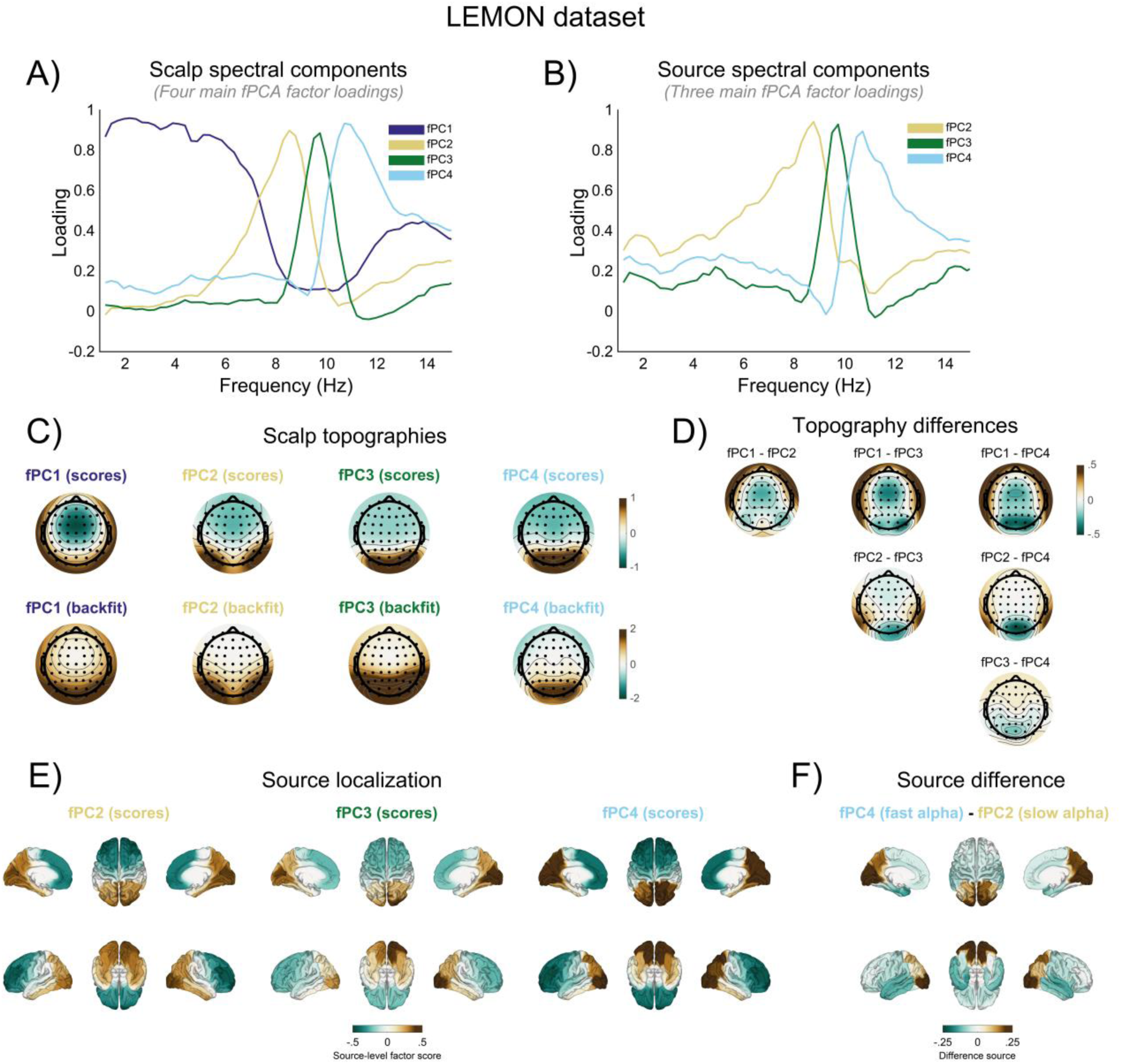
Alpha spectral components map onto distinct spatial and cortical patterns. A) In the LEMON dataset (N = 198), fPCA of scalp PSDs again identified four main spectral components explaining ≥80% of the variance, including three components within the alpha band. B) fPCA applied to cortical-source PSDs identified three main components explaining 87% of the variance, corresponding to the slow, middle, and fast alpha components. In panels A and B, results are displayed and color-coded as in Figures 1 and 2. C) Scalp topographies of component scores, obtained by averaging scores across participants at each electrode (top row), and back-fitted component topographies (bottom row; see Methods). D) Pairwise comparisons of component score topographies, showing the differences in scalp distribution across components. E) Source-level fPCA results projected onto 68 cortical regions (see Methods) recovered the three posterior alpha components and revealed distinct cortical expression patterns (scores estimated for each cortical region). F) Difference map between the slow and fast alpha component scores at the source level, highlighting relative differences in cortical expression. The fast alpha component showed stronger expression in dorsal occipito-parietal regions, whereas the slow alpha component was more expressed in lateral occipital and inferior temporal areas, suggesting a spatial dissociation broadly consistent with dorsal–ventral functional organization.

The LEMON dataset is multimodal and includes T1-weighted MRI for each participant. Using forward models derived from individual anatomy, we reconstructed EEG signals at the cortical level, estimated PSDs, and applied the same fPCA-based spectral decomposition used for the scalp data (see Methods). Using the same criterion of at least 5% explained variance, this analysis identified three alpha components in the source PSDs, together accounting for 87% of the variance (27.5% for fPC1, 42.5% for fPC2, and 17% for fPC3), but did not recover the low-frequency non-alpha component. The spectral profiles of these source-level components closely matched those obtained at the scalp (Figure 3B).

To characterize their spatial organization, we averaged component scores across cortical sources within the 68 regions defined by the Lausanne parcellation (scale 1; see Methods). All three components showed strong expression in occipito-parieto-temporal regions (Figure 3E). Importantly, the slow and fast alpha components exhibited distinct cortical distributions: the fast component showed higher expression in dorsal occipito-parietal areas, whereas the slow component showed higher expression in lateral occipital and inferior temporal regions (Figure 3F).

This dissociation confirms that the multiple alpha components identified at the scalp reflect partially distinct cortical generators.

## Discussion

For more than a century, alpha activity has been considered among the key biomarkers of brain function and dysfunction ^[51,3]^. One of the most extensively investigated features, the IAPF, has been used as a global measure to quantify the speed of alpha activity, implicitly assuming a continuum in which parametric variations relate to inter-individual variability in a range of perceptual and cognitive functions ^[12,52,53,9]^. Here, across multiple independent datasets, we show that the conventional estimate of IAPF does not reflect a unitary global rhythm. Instead, we show that IAPF emerges from the superposition of multiple alpha components with distinct spectral and spatial characteristics. These components are differentially expressed across individuals, and their relative contributions form stable individual profiles. Inter-individual variability in alpha activity is therefore better described as a compositional property rather than as parametric variation along a single-frequency continuum.

Our main finding is that highly similar spectral profiles and archetypal expression patterns can be consistently recovered across datasets that differ substantially in sample size, recording setup, laboratory of origin, and preprocessing pipelines. Across all datasets, the same set of components explained a large proportion of resting-state EEG variance, revealing that resting-state EEG is composed of a mixture of spectral components, including three robust alpha-band components. To our knowledge, this provides the first large-scale evidence that scalp EEG alpha activity is inherently compositional, and that its component structure is stable and reproducible at the population level.

These findings provide support for a body of previous work suggesting the coexistence of multiple alpha rhythms rather than a single global generator. Evidence for multiple alpha generators has been reported both at rest ^[28–30]^ and during active tasks ^[31–35]^, and was already hypothesized decades ago ^[27]^. Related ideas are also implicit in the long-standing practice of subdividing alpha activity into “lower” and “upper” bands ^[54,2]^, as well as in reports of individuals exhibiting multiple clearly identifiable alpha peaks in their power spectra ^[55]^. However, such distinctions have typically remained secondary and, at least in the context of estimating IAPF, have largely been overshadowed by the dominant practice of relying on a single global measure. Our results go beyond these earlier approaches by demonstrating that alpha activity is not merely divisible along an arbitrary frequency boundary, but instead reflects a structured mixture of reproducible components with distinct spectral properties, spatial organization, and stable individual expression patterns.

It must be noted that the approaches used here, fPCA and archetypal analysis, are designed to identify dominant directions of variance or extreme spectral profiles across an entire population. As such, they revealed canonical spectral components that characterize the main modes of variability within the datasets analyzed. Thus, they are not intended to provide an exhaustive description of all sources of individual variability and may not capture weaker or highly individual-specific alpha components that explain a smaller proportion of the variance. In line with this, our findings should not be interpreted as evidence that only three alpha generators exist. Rather, they indicate that a limited set of reproducible alpha components accounts for a substantial fraction of the population-level variability in resting-state EEG. Additional variability in the spectral shape, frequency, and spatial distribution of alpha activity is likely present at the individual level and may be better characterized using approaches specifically designed to capture subject-specific structure ^[56,57]^.

Furthermore, because the present analyses focused on eyes-closed resting-state EEG, the dominant alpha components were primarily expressed over posterior scalp regions, including occipital, parietal, and temporo-occipital electrodes. Different recording conditions may reveal additional alpha-band components with distinct functional and anatomical origins, such as sensorimotor mu and auditory-related tau rhythms ^[29,58,59]^, which occupy a similar frequency range but are typically maximal over central and temporal electrodes, respectively. Consistent with this possibility, applying the same spectral decomposition to eyes-open resting-state data (Supplementary Material “Spectral decomposition of eyes-open RS recording” and Figure S9) revealed two alpha components, corresponding to the slow and middle alpha components, rather than the three identified in eyes-closed data. The decomposition also recovered the non-alpha component and an additional component with loadings concentrated at higher frequencies. This pattern is consistent with previous findings from eyes-open resting-state EEG ^[60]^ and active tasks ^[61]^, suggesting that the four components identified in the present study reflect canonical spectral components of eyes-closed resting-state EEG rather than an exhaustive taxonomy of alpha-band activity. Nevertheless, given that many approaches rely on estimating a single IAPF from eyes-closed resting-state EEG, our findings highlight the clear limit and potential pitfalls of this single estimate.

An important consideration, however, is that the components identified by these approaches require careful interpretation, particularly when using fPCA. As demonstrated by our simulations (Figure S1), fPCA can separate a single underlying alpha component into multiple components when its peak frequency varies across individuals. Moreover, components identified at the population level may reflect distinct subgroups of individuals rather than multiple components coexisting within the same individual. Thus, the number of components recovered by a decomposition alone does not establish that they capture distinct sources of variability expressed within individuals. Their interpretation should instead rely on converging evidence, including the inspection and comparison of component score topographies, measures of component expression within individuals, and complementary decomposition approaches, such as archetypal analysis, which does not impose the same orthogonality constraints as fPCA (see also Supplementary Material, “Non-negative matrix factorization”). In the present study, we combined these approaches to ensure that the results were not driven by any specific decomposition method or its underlying assumptions.

If IAPF reflects the weighted summation of multiple underlying alpha components, then inferences drawn from the global IAPF alone are inherently ambiguous and, in some cases, potentially misleading. In particular, a shift in IAPF cannot be straightforwardly interpreted as a uniform speeding or slowing of alpha activity. Instead, it may reflect changes in the relative expression of distinct components, such as slower versus faster alpha components, without any change in their intrinsic frequencies. This fundamentally alters the interpretation of IAPF as a biomarker. Supporting this point, across the datasets analyzed, we show that distinct IAPF estimates can be obtained from electrode subsets preferentially expressing the slow and fast alpha components (see also Figures S5 and S8). These estimates capture partly non-overlapping information and may therefore differ in their associations with behavioral performance or clinical outcomes.

The association with age provides a clear example. Numerous studies have reported that IAPF follows a non-monotonic trajectory across the lifespan and that the slowing observed in older adults is associated with a general decline in cognitive abilities ^[48]^, leading to the interpretation of IAPF as a neurophysiological marker of global brain functioning ^[53]^. Our results suggest a different interpretation: age-related changes in IAPF primarily reflect shifts in the relative expression of specific alpha components rather than a homogeneous slowing of a single oscillatory process (see also ^[62]^). Our results also show that one of the alpha components (as well as the non-alpha component, Figure 1L) explains variance in age that is not captured by IAPF alone, resulting in an improved model (see also Figure S6). The age trajectories in the TDBRAIN dataset further suggest that the increase in IAPF typically observed during early development ^[63]^ involves both a decrease in the non-alpha component and an increase in the fast alpha component, with a smaller increase in the mid-alpha component (Figure 1L). That is, the developmental increase in IAPF may partly reflect an increasing contribution of faster alpha components. In contrast, the slow alpha component, which increases with older age, appears relatively stable during early development. This may indicate that distinct components and underlying neural circuits contribute to changes in the compositional structure of the EEG spectrum during early development and aging, the characterization of which remains an important question for future research.

A similar argument applies to schizophrenia. Previous studies have reported a slower IAPF in individuals with schizophrenia and have linked this effect to disease-related perceptual and cognitive deficits ^[64–67]^. However, prior work using the same spectral decomposition approach employed here has shown that schizophrenia is instead associated with the emergence or enhancement of a distinct lower-alpha/theta component with a specific spatial topography ^[43]^. Taken together, these findings clearly show how many results attributed to global changes in IAPF may instead arise from alterations in the compositional structure of alpha activity, reflecting not simply “slower” alpha activity, but a quantitatively different mixture of underlying generators.

As mentioned above, previous studies have already reported multiple alpha components with patterns largely consistent with those observed here ^[29,43,62,68–70]^. Building on this work, our large-scale investigation provides a general and decisive framework supporting these earlier findings, demonstrating that the compositional structure of alpha activity constitutes a highly stable and reliable neural trait. Importantly, we show that these components, and their relative contributions at the individual level, can be robustly estimated even from relatively short resting-state recordings and modest sample sizes, comparable to those commonly used in event-related EEG studies (Figure S8). This provides a practical tool for moving beyond brain-behavior associations based on a single, ambiguous global IAPF. This framework is also critical for causal intervention approaches, such as magnetic or electric brain stimulation, which aim to modulate ongoing alpha rhythms ^[71–75]^. Our results indicate that both stimulation target sites and stimulation frequency should be informed by the underlying compositional structure of alpha activity rather than by a unitary IAPF.

In general, we recommend that future studies investigating IAPF consider spectral decomposition approaches rather than relying exclusively on a single global estimate. Our findings, together with the shared code (see Code availability), provide a basis for evaluating different methods and metrics for this purpose. We recommend, for instance, combining metrics derived from fPCA and archetypal analysis as well as evaluating non-negative matrix factorization. However, given the consistency of the spectral loading profiles, scalp topographies, and contributing electrodes associated with each component, simpler approaches may also be useful at the individual level. These include: (1) back-projecting canonical spectral component topographies derived from an independent dataset onto individual data, as illustrated in Figure 3C, in a manner analogous to microstate analysis ^[76]^; and (2) jointly assessing associations with IAPF and variables of interest by estimating IAPF from electrode subsets preferentially expressing different components, as identified here, after verifying that the resulting estimates capture partly non-overlapping information.

A key question naturally raised by our findings is about the functional roles, if any, to which distinct alpha components can be mapped. In source analysis, we found that the two components most strongly involved in explaining variability in both IAPF and age were associated with sources located in occipito-parietal (fast alpha) and occipito-temporal (slow alpha) regions. These source locations are consistent with distinct alpha generators reported in previous studies using smaller samples ^[29]^, and have recently been linked to different functional roles ^[77]^. Their cortical distribution is also loosely reminiscent of the classic distinction between dorsal and ventral visual pathways, suggesting that different alpha components may support partially dissociable computational or functional roles, potentially related to distinct streams of visuo-spatial processing, a hypothesis with important implications for future research. It should be noted that while scalp-level fPCA systematically revealed an additional low-frequency, non-alpha component, this component did not meet the variance-based selection criterion in source space (5% explained variance), accounting instead for 4.6% of the variance. One possibility is that this component is attenuated in source space because it reflects more spatially diffuse, low signal-to-noise activity. In contrast, alpha components remain spatially coherent and continue to dominate the low-dimensional structure of source-level spectra.

We hope that focusing on the compositional nature of alpha activity, rather than on the global IAPF, can provide a more fine-grained tool to help resolve inconsistent findings in the literature and replication failures ^[78–82]^. Essentially, we might have been looking at the wrong measure: true associations between behavioral metrics, cognitive scores, or clinical phenotypes and alpha components may have been conflated or weakened because these components were combined into a single global IAPF estimate. This is also highly relevant for clinical applications, where IAPF has been used, for example, to predict treatment ^[83,84]^, pain sensitivity ^[85]^, and for patient stratification ^[86]^. Similarly, from a methodological point of view, there are currently widespread methods to parametrize spectral components and estimate IAPF or other properties of oscillatory and “aperiodic” components in neural activity. These approaches, however, generally assume that spectral peaks reflect individual oscillatory processes superimposed on an aperiodic background. They also require several analytical choices ^[87–90]^ and often rely on fitting parameterized models to the log-transformed power spectrum, residualizing periodic peaks from the aperiodic background. Our approach and findings show instead that distinct alpha components, alongside lower-frequency and 1/f-like components, can be identified and separated in an unsupervised, and data-driven manner, allowing their expression to be quantified at both the scalp and individual-participant levels. These measures can then be readily incorporated into brain–behavior models and may provide more precise alpha-based biomarkers for clinical research. Future research should also assess whether, and to what extent, the lower-frequency non-alpha component identified here is related to the increasingly studied aperiodic component of the EEG power spectrum.

In sum, human alpha activity is composed of multiple rhythms with distinct scalp patterns that form a set of canonical eyes-closed resting-state EEG alpha components. These components combine in different proportions within each individual, giving rise to a highly stable neural trait. Despite over a century of research, the functional roles of these components and their generating mechanisms remain largely unknown, obscured by the traditional reliance on a single IAPF estimate. We propose that compositional measures of alpha activity should be used to revisit the extensive literature on alpha rhythms, opening new avenues for more precise and mechanistic accounts of individual differences in brain function, as well as for the development of more informative alpha-based biomarkers.

## Material and methods

### Sample description

We used three publicly available large-scale datasets of resting-state EEG (RS-EEG) recordings, comprising a total of 2110 participants (see Supplementary Material and Figure S8 for validation in an additional, smaller (N = 19) in-house dataset ^[91]^).

The first dataset consists of RS-EEG recordings from 1299 participants, representing a heterogeneous sample of healthy individuals and patients with various clinical diagnoses, with ages ranging from 5 to 83 years (608 females, 691 males). These recordings were collected in Nijmegen, the Netherlands, over a 20-year period as part of the *Two Decades–Brainclinics Research Archive for Insights in Neurophysiology* (TDBRAIN; for details, see ^[39]^). Each participant completed a 4-minute RS-EEG recording, beginning with 2 minutes in an eyes-open (EO) condition and followed by 2 minutes in an eyes-closed (EC) condition.

The second dataset involves RS-EEG recordings from 608 healthy participants aged 20 to 70 years (376 females, 232 males), collected in Dortmund, Germany, as part of the *Dortmund Vital Study* (DVS; for details, see ^[49]^. For each participant, a 6-minute RS-EEG was initially recorded, with 3 minutes in an EC condition followed by 3 minutes in an EO condition (referred to as “pre-test” recordings). The participants then performed a series of five cognitive tasks lasting approximately two hours in total before completing a second 6-minute RS-EEG recording, again consisting of 3 minutes in both EC and EO conditions (referred to as “post-test” recordings). A subset of 208 participants (130 females, 78 males) returned for a second session, approximately five years later, following the same protocol.

The third dataset contains RS-EEG recordings from 203 healthy participants, collected in Leipzig, Germany, as part of the *Leipzig Study for Mind-Brain-Body Interactions* (LEMON; for details, see ^[50]^. Participants were divided into two age groups: 138 young adults aged 20 to 35 years (42 females, 96 males) and 65 older adults aged 59 to 77 years (32 females, 33 males). Each participant completed a 16-minute resting-state EEG recording with 16 alternating 60-second blocks of EC and EO conditions, always starting with an EC block. In addition, individual structural magnetic resonance imaging (MRI) data were acquired for each of these participants (see ^[50]^ for details).

Data collection for the TDBRAIN, DVS, and LEMON datasets was conducted in accordance with local ethics committee guidelines and the Declaration of Helsinki. All participants provided written informed consent before data collection. For additional details, please refer to the original studies ^[50,39,49].^

### RS-EEG acquisition and preprocessing

RS-EEG recordings from the TDBRAIN dataset were obtained using either a Quickcap (Compumedics, NC, USA) or an ANT-Neuro Waveguard Cap (Brain Products GmbH, Gilching, Germany) equipped with sintered Ag/AgCl electrodes. The EEG setup comprised 26 channels arranged according to the international 10–10 system. Signals were recorded at a sampling rate of 500 Hz. A virtual ground was used during acquisition, with offline re-referencing to the averaged mastoids (A1 and A2) and a ground placed at AFz. Skin–electrode impedance was maintained below 10 kΩ throughout the recordings.

For our study, we preprocessed the 2-minute EC segments from the TDBRAIN dataset using an automated pipeline recommended by Delorme (2023) in EEGLAB (version v2024.2 ^[93]^). First, EEG recordings were downsampled from 500 Hz to 250 Hz and a 0.5 Hz high-pass Finite Impulse Response (FIR) filter (*pop_eegfiltnew()*) was applied. Next, we identified bad channels and noisy data segments using the *clean_rawdata* plugin: Channels with correlation values below 0.85 and data segments exceeding the Artifact Subspace Reconstruction (ASR ^[94]^) threshold of 20 were removed. Independent component analysis (ICA) was then performed using Infomax (*Picard* plugin, *pop_runica()*) and components associated with muscle and eye artifacts, identified with >90% probability by the *ICLabel* algorithm, were removed. After preprocessing, an average of 1.9 electrodes, 13% of the data, and 0.7 independent components were discarded across the 2 minutes of EC recordings. Finally, removed electrodes were interpolated using spherical spline interpolation (*pop_interp()* ^[95]^), all electrodes were re-referenced to the average reference (*pop_reref()*), and the clean data were epoched into 2-second windows. To ensure a minimal amount of data per participant, we excluded 19 participants for whom the automated pipeline retained less than 1 minute of recording, resulting in 1270 retained participants from the initial 1299. The same preprocessing pipeline was applied to the EO recordings (see Supplementary Material and Figure S9).

RS-EEG data from the DVS dataset were recorded with a BrainVision BrainAmp DC amplifier using a 64-channel elastic cap (Brain Products GmbH, Gilching, Germany), configured according to the 10–20 system, with the FCz electrode serving as the online reference. The EEG signal was sampled at 1000 Hz and all electrode impedances were maintained below 10 kΩ.

From the DVS dataset, we considered and preprocessed the 3-minute EC segments, recorded before and after the cognitive tasks, during the first session and, when available, the 3-minute EC segments from the second session 5 years later. We followed the same automated pipeline, as described above for the TDBRAIN dataset, except that EEG recordings were downsampled here from 1000 Hz to 250 Hz. An average of 3.5 electrodes, 15.5% of the data, and 4.7 independent components were discarded across the 12 minutes of EC recordings. Finally, participants were excluded if the automated pipeline retained less than 1 min of usable data in either the pre- or post-test EC recordings within a given session. Of the 608 participants in Session 1, 604 were retained for the pre-test recordings (4 excluded) and 605 for the post-test recordings (3 excluded). Similarly, of the 208 participants in Session 2, 207 were retained for the pre-test recordings (1 excluded) and 206 for the post-test recordings (2 excluded). The 3-minute EO segments from both sessions (pre-test) were also preprocessed (see Supplementary Material and Figure S9).

RS-EEG data from the LEMON dataset were recorded with a BrainVision BrainAmp MR plus amplifier using 61 active ActiCAP electrodes (Brain Products GmbH, Gilching, Germany), arranged according to the 10-10 international system, with the FCz electrode serving as the online reference. The ground electrode was positioned on the sternum, and all electrode impedances were kept below 5 kΩ. Data were continuously recorded with an online bandpass filter ranging from 0.015 Hz to 1 kHz and digitized at a sampling rate of 2500 Hz.

We used the EEG data preprocessed by Babayan and colleagues ^[50]^. Specifically, after data acquisition, EEG recordings were downsampled from 2500 Hz to 250 Hz, bandpass filtered between 1 and 45 Hz using an eighth-order Butterworth filter, and separated into EO and EC conditions. Next, electrodes exhibiting frequent voltage shifts or poor signal quality were identified via visual inspection and excluded. Similarly, data segments containing extreme peak-to-peak deflections or high-frequency bursts were manually removed. Dimensionality reduction was then performed using principal component analysis (PCA), retaining a sufficient number of principal components (N ≥ 30) to explain 95% of the total variance and independent component analysis using Infomax (*Picard* plugin, *pop_runica()*) was applied to visually identify and remove components associated with eye movements, blinks, or cardiac activity. Further information about EEG data acquisition and preprocessing can be found in Babayan et al. ^[50]^. In the present work, we applied additional preprocessing steps that included interpolation of removed electrodes, re-referencing to the average reference, and epoching into 4-second windows. In the EC recordings, two participants were excluded due to differing sampling rates in their original EEG recordings, resulting in 201 retained participants (138 young adults and 63 older adults; see Supplementary Material and Figure S9 for EO recordings).

Although the automated preprocessing pipelines for the TDBRAIN and DVS datasets are similar, substantial differences in sample size (1270 vs. 604 participants retained after preprocessing) and the number of electrodes (26 vs. 64) provide additional control, ensuring that our results are not driven solely by sample size or EEG coverage. The same applies to the LEMON dataset (201 participants retained; 61 electrodes) and an additional in-house dataset (19 participants; 128 electrodes, see Supplementary Material and Figure S8), which were additionally preprocessed using different pipelines and segmented into longer epoch durations (4 seconds vs. 2 seconds in TDBRAIN and DVS). These differences further reduce the likelihood that our findings are specific to, or artifacts of, a particular preprocessing pipeline.

### Power spectral density

For each participant in all datasets, we estimated the power spectral density (PSD) at each electrode and epoch within the 1–40 Hz frequency range using a 1024-point fast Fourier transform (*fft*(), MATLAB R2024b, frequency resolution of 0.2441 Hz). Prior to PSD estimation, resting-state EEG (RS-EEG) data were globally z-scored across all electrodes and epochs, and single epochs were demeaned.

To estimate the individual alpha peak frequency (IAPF), we employed the FOOOF algorithm, which separates the 1/f background component from oscillatory peaks in the PSD ^[89]^. FOOOF was applied to the PSD averaged across electrodes and epochs, with the most prominent peak in the 7–14 Hz range identified as the IAPF. Default parameters were used (peak_width_limits = (0.5, 12.0), max_n_peaks = inf, min_peak_height = 0, peak_threshold = 2.0, aperiodic_mode = ‘fixed’). Participants for whom IAPF detection via FOOOF was unsuccessful were excluded from further analysis. This resulted in final sample sizes of 1268 participants from the TDBRAIN dataset (12 excluded); 595 and 601 participants for the pre- and post-test recordings from the first session in the DVS dataset (9 and 4 excluded, respectively); 206 participants for the pre- and post-test recordings from the second session in the DVS dataset (1 and 0 excluded, respectively); 198 participants from the LEMON dataset (3 excluded); and 19 participants from the in-house dataset (0 excluded). Across all datasets, the average model fit quality was higher than 0.99.

### Spectral decomposition via frequency-PCA

To estimate the spectral components characteristics of RS-EEG we applied the frequency-PCA (frequency principal component analysis, fPCA) method proposed by Nakhnikian and colleagues ^[43]^ (see also ^[41,42]^). First, the PSD estimated at each electrode and for each participant, averaged over epochs, was concatenated in a matrix:

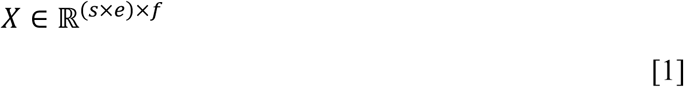

where *s* is the number of subjects, *e* is the number of electrodes and *f* is the number of frequency bins obtained from the PSD estimation (159 bins, restricted to the 1-40 Hz range). Each row of *X* corresponds to the PSD of a specific subject-electrode pair across frequencies. We then applied PCA:

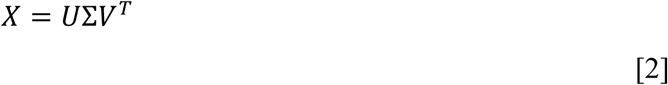

where *U* ∈ ℝ^(*s*×*e*)×(*s*×*e*)^, Σ ∈ ℝ^(*s*×*e*)×*f*^ is the diagonal matrix of singular values, and V ∈ ℝ*^f^*^×*f*^ contains the principal components. The columns of *V* correspond to the factor loadings, representing spectral profiles across frequencies. The matrix *U*Σ contains the factor scores, i.e. the expression of each spectral component for every subject–electrode observation. Thus, the first few components in *V* describe the dominant spectral modes present in the data, whereas the associated scores indicate how strongly each spectral component is expressed across electrodes and participants.

A PCA was initially conducted to determine the number of components to retain, using the criterion of retaining components that explained at least 5% of the variance in the PSD. This resulted in the retention of four components in all analyses performed (e.g., Figure 1D), except for source reconstruction (Figure 3B), which combined always accounted for more than 80% of the variance.

After performing a subsequent PCA with the retained components, we applied Varimax rotation following the approach in Nakhnikian and colleagues ^[43]^, i.e., using the PCA toolbox ^[96]^ (*doPCA*(), configured for covariance-based PCA with unscaled Varimax rotation and without Kaiser normalization). This orthogonal rotation maximizes the separation of components by simplifying their structure, such that each component loads strongly on a subset of frequencies and weakly on others. This improves interpretability of the resulting factor loadings in the original PSD space (e.g., Figure 1D). Spatial expressions of each spectral component were derived from the factor scores obtained during fPCA. Scores were reshaped into *s* × *e* × *k* arrays (where *k* indexes component), and scalp maps were obtained by averaging scores across participants for each electrode and component (e.g., Figure 3C). These maps represent the spatial distribution of component expression (factor scores) across the scalp and were computed at the single-subject level to provide a global measure of the contribution of each component to an individual’s scalp PSD, which was then used for predictive models (e.g., Figure 1L).

Unlike classic global IAPF approaches, which typically collapse spectral information across electrodes or rely on a predefined subset, the present method explicitly models the joint spectral– spatial structure of RS-EEG. That is, by decomposing PSDs at the scalp level, fPCA allows multiple alpha-related components with distinct spectral profiles and spatial distributions to be identified simultaneously. The resulting component score topographies therefore provide a key dimension for assessing the extent to which the identified components are spatially distinct, and thus whether the decomposition effectively separates different components, as further illustrated by simulations and within-individual electrode subtyping analyses in the Supplementary Material (Figures S1 and S3).

Additionally, to avoid potential confounds in the interpretation of component score topographies arising from PCA sign ambiguity, we also estimated component topographies using a back-fitting procedure. In this procedure, group-level spectral loading profiles were fitted to each participant’s PSD at each electrode using multiple linear regression. The regression coefficients associated with each component, excluding the intercept, were then used to construct participant-specific scalp maps of component expression (e.g., Figure 3C).

### Archetypal analysis

To further characterize spectral profiles at the group level, we applied archetypal analysis, implemented via Principal Convex Hull Analysis (*PCHA*() ^[44]^), to RS-EEG PSD. Unlike PCA, which identifies orthogonal components that maximize variance, archetypal analysis represents data as convex combinations of a small set of extreme patterns (“archetypes”), which lie on the boundary of the data cloud and can be interpreted as archetypal spectral profiles.

For this analysis, PSD estimates were first averaged across electrodes for each participant, yielding a subject-by-frequency matrix *X* ∈ ℝ*^s^*^×*f*^, where *s* denotes the number of subjects and *f*the number of frequency bins. To ensure non-negativity and comparability across participants, PSD values were rescaled to the [0,1] interval prior to analysis. Archetypal analysis was then performed on the transposed matrix *X*^T^, such that each archetype corresponds to a spectral profile across frequencies.

Formally, this analysis approximates the data matrix as:

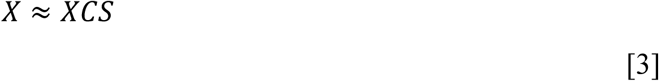

where the columns of *XC* define the archetypes, *C* is a non-negative matrix whose columns sum to one and select convex combinations of observed spectra to form archetypes, and *S* is a non-negative coefficient matrix whose columns sum to one and describe how strongly each archetype contributes to each participant’s spectrum. This formulation ensures that both archetypes and subject-level reconstructions remain within the convex hull of the observed data.

The number of archetypes *k* was selected in a data-driven manner by evaluating models with *k* = 2 to 10 archetypes and computing the fraction of variance explained by each solution. The smallest *k* accounting for more than 95% of the total variance was retained for subsequent analyses. As in the fPCA approach, this resulted in *k* = 4—i.e., four main archetypes. Final model estimation was performed using this value of *k*. Note that, in contrast to fPCA, which captures dominant modes of variance, archetypal analysis emphasizes extreme spectral patterns and represents individual participants as mixtures of these archetypal profiles. Nevertheless, both approaches converged to the identification of four main components/archetypes (e.g., Figure 1E). For consistency and graphical purposes, we reordered the archetypes based on their close correspondence with the spectral components identified via fPCA.

To characterize how strongly individual participants expressed specific archetypal spectral profiles, we quantified the *dominance* and *participation ratio* of archetypal representations based on the subject-level weight matrix *S*. Each column of *S* contains non-negative weights summing to one and describes the convex combination of archetypes used to reconstruct a given participant’s scalp-averaged PSD. We identified the dominant archetypes by taking the index of the maximum value, per participant, in the related row of *S*. We assessed the distribution of dominant archetypes across participants by computing the proportion of participants for whom each archetype had the highest weight (e.g., Figure 1G). This analysis provides a view of how frequently each archetypal spectral profile serves as the primary contributor to individual PSDs. The participation ratio (PR) was computed as 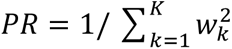 , where *w*_k_ denotes the normalized weight of each archetype, providing an estimate of the effective number of contributing archetypes. Profiles were classified as *pure* (PR < 1.5), *dual* (1.5 ≤ PR < 2.5), or *distributed* (PR ≥ 2.5), using thresholds to distinguish profiles dominated by a single archetype, those reflecting contributions from approximately two archetypes, and those with more broadly distributed contributions, respectively (e.g., Figure 1H).

To visualize how participants are distributed in the reduced spectral space of components/archetypes, we projected individual archetypal expressions into the three alpha-related fPCs obtained from the fPCA scalp-averaged scores and represented each participant as a point in a three-dimensional fPC space (e.g., Figure 1F). Each point therefore corresponds to a participant’s position along the dominant spectral component axes, capturing the major sources of inter-individual spectral variability as derived from fPCA. Participants were color-coded according to their dominant archetype, defined as the archetype with the highest weight for that individual. The size of each point was scaled according to its participation ratio, with larger points indicating a more distributed contribution across archetypes. This visualization allows a qualitative assessment of how archetypal dominance relates to the geometry of the spectral space, that is, whether participants associated with different archetypes occupy distinct regions, form clusters, or instead exhibit smooth transitions across the fPC dimensions.

### Relationships between IAPF and the spectral components

We assessed the relationship between IAPF and the spectral components in the TDBRAIN dataset. A multiple linear regression model was fitted with IAPF as the dependent variable and the scalp-averaged scores associated with the four spectral components identified by the fPCA as predictors. For this model, we reported regression coefficients (β), standard errors, test statistics, and *p* values for each predictor. Given the strong and opposing contributions of the two dominant alpha-related components (corresponding to slow and fast alpha; e.g., Figure 1J), we conducted an additional analysis to quantify their shared relationship with IAPF. A contrast term was computed as the difference between the slow- and fast-alpha component scores, and a separate linear regression model was fitted with this contrast as the sole predictor of IAPF.

In addition, to assess whether slow and fast alpha components provide distinct information about IAPF, we identified electrode subsets showing preferential expression of each component based on the group-level component score topographies. For each electrode, selectivity was quantified as the difference between the spatial scores of the slow and fast alpha components, and the five electrodes with the highest selectivity scores were selected for each subset. IAPF was then estimated separately from the PSDs averaged across each electrode subset using FOOOF. The degree of shared variance between the two IAPF estimates was quantified using the R^2^ from a linear regression, with lower values indicating less shared variance (e.g., Figure 1K).

Similar analyses were also conducted using the DVS and LEMON datasets (see Figure S5).

### Age prediction

IAPF is known to vary systematically across the lifespan, increasing during early development and declining in later adulthood ^[39,38]^. As an initial validation step, we confirmed this well-established relationship in the TDBRAIN dataset, via linear and quadratic regression models fitted with IAPF as the dependent variable and age as the predictor (Figure 1L). Model selection (e.g., linear vs. quadratic) was performed using the Bayesian Information Criterion (BIC), and the model with the lower BIC was retained. For the selected model, regression coefficients (β), standard errors, test statistics, and associated *p* values for the age terms were reported.

To further determine whether distinct age-related changes were present in each spectral component identified by the fPCA analysis in the TDBRAIN dataset, linear and quadratic regression models were also fitted with the scalp-averaged score associated with each spectral component as the dependent variable and age as the predictor (e.g., Figure 1L). Model selection was again based on BIC, with the model exhibiting the lower BIC retained. For each selected model, regression coefficients (β), standard errors, test statistics, and p values for the age terms were reported. Adjusted R² values were also reported to characterize the proportion of age-related variance explained by each component. Similar models were fitted to the DVS and LEMON datasets (see Figure S6).

### Stability and fingerprinting analysis

As mentioned, the DVS dataset included multiple RS-EEG recordings from the same individuals, with two recordings on the same day (i.e., pre- and post-test recordings from the first session) and, for a subset of participants, two additional recordings approximately five years later (i.e., pre- and post-test recordings from the second session). This longitudinal design allowed us to evaluate the stability of spectral component scores across both short- and long-term intervals. Scores derived from the fPCA decomposition were averaged across electrodes for each participant and session. Within-session test-retest stability was quantified by correlating these scalp-averaged scores between pre-and post-test recordings on the same day. We used linear regression to model the scores in one session as a function of those in the other, and we assessed the adjusted R² values as a measure of the reliability of each spectral component (Figure 2E). Longitudinal stability across sessions separated by several years was assessed in the same manner (Figure 2D). Only participants with data available for both recordings in a given comparison were included, resulting in 590 participants for pre-vs post-test recordings from the first session and 202 participants for pre-test recordings from the first session vs pre-test recordings from the second session.

To assess the uniqueness of individual spectral profiles, we performed a fingerprint analysis ^[97,98]^. We computed a subject-by-subject correlation matrix on vectorized topographies of component scores across two sessions, both within the same day, and after five years. That is, we correlated vectors obtained by concatenating electrode-by-component scores, capturing the full spatial expression of all components for each participant, between sessions. The diagonal elements of this correlation matrix (ISelf) quantify, for each participant, the correlation between their own scalp spectral profile across sessions. In contrast, the mean correlation between a participant’s profile and those of all other participants (IOthers) captures between-subject similarity. The difference between the population means of ISelf and IOthers (IDiff) provides an estimate of subject-specific reliability, while the proportion of participants for whom ISelf exceeds IOthers defines the identifiability success rate (Figure 2F). High IDiff values and high success rates indicate that individual spectral profiles are both stable over time and distinguishable across participants, supporting the use of spectral component factor scores as reliable individual “fingerprint”.

### EEG inverse modeling

The LEMON dataset includes individual structural magnetic resonance imaging (MRI) data acquired from the same participants who underwent RS-EEG recordings (see ^[50]^ for detailed acquisition parameters). Anatomical T1-weighted MRI data were preprocessed using the Connectome Mapper 3 (v3.1.0; ^[99]^) open-source pipeline with FreeSurfer (v7.1.1) software package. Because digitized electrode positions were not available for all participants, we employed a standard 64-channel BioSemi EEG cap template. This template montage was co-registered to a template anatomical MRI in MNI space, and then warped to each individual’s native space using the nonlinear transformation matrix from MNI to individual anatomy.

T1-weighted MRI data were segmented to extract scalp, skull, and brain surfaces. Volume conduction models were constructed using the boundary element method (BEM) implemented in OpenMEEG ^[100]^. Source models were defined using a regular 7 mm grid of equivalent current dipoles constrained to the cortical gray matter of the MNI template brain, yielding 2501 solution points in cortical tissue. Each participant’s anatomical image was then registered to the MNI template, and the inverse deformation field was applied to project the MNI grid into individual anatomical space ^[101,102]^.

Forward models were computed for each participant using unconstrained dipole orientations, allowing three orthogonal dipole components per location. EEG source activity was reconstructed using standardized low-resolution brain electromagnetic tomography (sLORETA) ^[103]^ with a regularization parameter λ = 10% . For each dipole and direction, the PSD was computed per epoch, then averaged across the three dipole orientations at each solution point using the same method as in the scalp-level analysis.

Single-subject PSD data from all solution points were concatenated and analyzed using the fPCA approach described above, replacing electrode-level input with source-level data. Components were retained using the same criterion of explaining at least 5% of variance, which in the source analysis yielded three components corresponding to the three alpha-related sources (Figure 3B), excluding the low-frequency non-alpha component identified at the scalp (e.g., Figure 3A). Seven participants were excluded from the source-level analysis due to failed anatomical segmentation or forward model computation, likely caused by the facial anonymization procedure in the LEMON dataset that affected tissue segmentation and head model accuracy. For visualization, component scores were averaged within 68 cortical regions of interest (Figure 3E) defined by the Lausanne parcellation (scale 1) ^[104]^, which is derived from the Desikan–Killiany anatomical atlas ^[105]^. Only cortical gray-matter regions were included. In addition, to assess whether the slow and fast alpha components exhibited distinct cortical source distributions, we computed a difference map by subtracting the slow alpha component scores from the fast alpha component scores across cortical regions (Figure 3F).

## Acknowledgments

This work was supported by the Swiss National Science Foundation (Grant number: TMSGI1_218247). We thank the authors of the datasets used in this study for making these valuable resources openly available, and Songyun Bai for support with the collection of the in-house dataset.

## Author contribution

Conceptualization: MQM & DP

Methodology: MQM & DP

Investigation: MQM & DP

Visualization: MQM & DP

Supervision: DP

Writing—original draft: MQM & DP

Writing—review & editing: MQM & DP

## Code availability

The code used in this study to analyze EEG power spectra, including PSD estimation and preprocessing, fPCA, archetypal analysis, NMF, simulation routines, and utility functions, is available in an online repository (DOI: https://doi.org/10.5281/zenodo.23600677).

## Data availability

The datasets analyzed in this study are publicly available through the repositories linked in their original publications (TDBRAIN dataset, https://www.synapse.org/Synapse:syn25671079/wiki/610278; DVS dataset, https://openneuro.org/datasets/ds005385/versions/1.0.0; LEMON dataset, https://openneuro.org/datasets/ds000221/versions/1.0.0). The in-house sample dataset used toillustrate the application of the code is available in an online repository (DOI: https://doi.org/10.5281/zenodo.23600677).

## References

1. Berger, H. (1929). Über das elektroenkephalogramm des menschen. Archiv für psychiatrie und nervenkrankheiten, 87(1), 527–570.

2. Klimesch, W. (1999). EEG alpha and theta oscillations reflect cognitive and memory performance: A review and analysis. Brain Research Reviews, 29(2–3), 169–195. 10.1016/S0165-0173(98)00056-3

3. Pascucci, D., Menétrey, M. Q., Passarotto, E., Luo, J., Paramento, M., & Rubega, M. (2025). EEG brain waves and alpha rhythms: Past, current and future direction. Neuroscience & Biobehavioral Reviews, 176, 106288. 10.1016/j.neubiorev.2025.106288

4. Haegens, S., Cousijn, H., Wallis, G., Harrison, P. J., & Nobre, A. C. (2014). Inter- and intra-individual variability in alpha peak frequency. NeuroImage, 92, 46–55. 10.1016/j.neuroimage.2014.01.049

5. Grandy, T. H., Werkle-Bergner, M., Chicherio, C., Schmiedek, F., Lövdén, M., & Lindenberger, U. (2013). Peak individual alpha frequency qualifies as a stable neurophysiological trait marker in healthy younger and older adults: Alpha stability. Psychophysiology, 50(6), 570–582. 10.1111/psyp.12043

6. Popov, T., Tröndle, M., Baranczuk-Turska, Z., Pfeiffer, C., Haufe, S., & Langer, N. (2023). Test–retest reliability of resting-state EEG in young and older adults. Psychophysiology, 60(7), e14268. 10.1111/psyp.14268

7. Arioli, M., Mattersberger, M., Hoehl, S., & Brzozowska, A. (2024). Peak alpha frequency is linked to visual temporal attention in 6-month-olds. Scientific Reports, 14(1), 28173. 10.1038/s41598-024-79129-0

8. Menétrey, M. Q., Roinishvili, M., Chkonia, E., Herzog, M. H., & Pascucci, D. (2024). Alpha peak frequency affects visual performance beyond temporal resolution. Imaging Neuroscience, 2, 1–12. 10.1162/imag_a_00107

9. Samaha, J., & Romei, V. (2024). Alpha-Band Frequency and Temporal Windows in Perception: A Review and Living Meta-analysis of 27 Experiments (and Counting). Journal of Cognitive Neuroscience, 36(4), 640–654. 10.1162/jocn_a_02069

10. Marsicano, G., Deodato, M., & Melcher, D. (2026). Alpha oscillations and aperiodic neural dynamics jointly predict visual temporal resolution, confidence, and dependence on prior experience. 10.7554/eLife.110300.1

11. Ramsay, I. S., Lynn, P. A., Schermitzler, B., & Sponheim, S. R. (2021). Individual alpha peak frequency is slower in schizophrenia and related to deficits in visual perception and cognition. Scientific Reports, 11(1), 17852. 10.1038/s41598-021-97303-6

12. Klimesch, W., Schimke, H., & Pfurtscheller, G. (1993). Alpha frequency, cognitive load and memory performance. Brain Topography, 5(3), 241–251. 10.1007/BF01128991

13. Ociepka, M., Kałamała, P., & Chuderski, A. (2022). High individual alpha frequency brains run fast, but it does not make them smart. Intelligence, 92, 101644. 10.1016/j.intell.2022.101644

14. Babu Henry Samuel, I., Wang, C., Hu, Z., & Ding, M. (2018). The frequency of alpha oscillations: Task-dependent modulation and its functional significance. NeuroImage, 183, 897–906. 10.1016/j.neuroimage.2018.08.063

15. Grandy, T. H., Werkle-Bergner, M., Chicherio, C., Lövdén, M., Schmiedek, F., & Lindenberger, U. (2013). Individual alpha peak frequency is related to latent factors of general cognitive abilities. NeuroImage, 79, 10–18. 10.1016/j.neuroimage.2013.04.059

16. Ippolito, G., Bertaccini, R., Tarasi, L., Di Gregorio, F., Trajkovic, J., Battaglia, S., & Romei, V. (2022). The Role of Alpha Oscillations among the Main Neuropsychiatric Disorders in the Adult and Developing Human Brain: Evidence from the Last 10 Years of Research. Biomedicines, 10(12), 3189. 10.3390/biomedicines10123189

17. Eidelman-Rothman, M., Levy, J., & Feldman, R. (2016). Alpha oscillations and their impairment in affective and post-traumatic stress disorders. Neuroscience & Biobehavioral Reviews, 68, 794–815. 10.1016/j.neubiorev.2016.07.005

18. Finley, A. J., Angus, D. J., Knight, E. L., Van Reekum, C. M., Lachman, M. E., Davidson, R. J., & Schaefer, S. M. (2024). Resting EEG Periodic and Aperiodic Components Predict Cognitive Decline Over 10 Years. The Journal of Neuroscience, 44(13), e1332232024. 10.1523/JNEUROSCI.1332-23.2024

19. Lejko, N., Larabi, D. I., Herrmann, C. S., Aleman, A., & Ćurčić-Blake, B. (2020). Alpha Power and Functional Connectivity in Cognitive Decline: A Systematic Review and Meta-Analysis. Journal of Alzheimer’s Disease, 78(3), 1047–1088. 10.3233/JAD-200962

20. Janiukstyte, V., Kozma, C., Owen, T. W., Chaudhary, U. J., Diehl, B., Lemieux, L., Duncan, J. S., Rugg-Gunn, F., De Tisi, J., Wang, Y., & Taylor, P. N. (2024). Alpha rhythm slowing in temporal lobe epilepsy across scalp EEG and MEG. Brain Communications, 6(6), fcae439. 10.1093/braincomms/fcae439

21. Dickinson, A., DiStefano, C., Senturk, D., & Jeste, S. S. (2017). Peak alpha frequency is a neural marker of cognitive function across the autism spectrum. European Journal of Neuroscience, 9.

22. Babiloni, C., Jakhar, D., Tucci, F., Del Percio, C., Lopez, S., Soricelli, A., Salvatore, M., Ferri, R., Catania, V., Massa, F., Arnaldi, D., Famà, F., Güntekin, B., Yener, G., Stocchi, F., Vacca, L., Marizzoni, M., Giubilei, F., Yıldırım, E., … Noce, G. (2024). Resting state electroencephalographic alpha rhythms are sensitive to Alzheimer’s disease mild cognitive impairment progression at a 6-month follow-up. Neurobiology of Aging, 137, 19–37. 10.1016/j.neurobiolaging.2024.01.013

23. Cecchetti, G., Agosta, F., Canu, E., Basaia, S., Rugarli, G., Curti, D. G., Coraglia, F., Cursi, M., Spinelli, E. G., Santangelo, R., Caso, F., Fanelli, G. F., Magnani, G., & Filippi, M. (2024). Analysis of individual alpha frequency in a large cohort from a tertiary memory center. European Journal of Neurology, 31(10). 10.1111/ene.16424

24. Zhao, Y., Cai, J., Song, J., Shi, H., Kong, W., Li, X., Wei, W., & Xue, X. (2025). Peak alpha frequency and alpha power spectral density as vulnerability markers of cognitive impairment in Parkinson’s disease: An exploratory EEG study. Frontiers in Neuroscience, 19, 1575815. 10.3389/fnins.2025.1575815

25. Murphy, M., & Öngür, D. (2019). Decreased peak alpha frequency and impaired visual evoked potentials in first episode psychosis. NeuroImage: Clinical, 22, 101693. 10.1016/j.nicl.2019.101693

26. Zhou, P., Wu, Q., Zhan, L., Guo, Z., Wang, C., Wang, S., Yang, Q., Lin, J., Zhang, F., Liu, L., Lin, D., Fu, W., & Wu, X. (2023). Alpha peak activity in resting-state EEG is associated with depressive score. Frontiers in Neuroscience, 17, 1057908. 10.3389/fnins.2023.1057908

27. Başar, E., Schürmann, M., Başar-Eroglu, C., & Karakaş, S. (1997). Alpha oscillations in brain functioning: An integrative theory. International Journal of Psychophysiology, 26(1–3), 5–29. 10.1016/S0167-8760(97)00753-8

28. Ben-Simon, E., Podlipsky, I., Arieli, A., Zhdanov, A., & Hendler, T. (2008). Never Resting Brain: Simultaneous Representation of Two Alpha Related Processes in Humans. PLoS ONE, 3(12), e3984. 10.1371/journal.pone.0003984

29. Barzegaran, E., Vildavski, V. Y., & Knyazeva, M. G. (2017). Fine Structure of Posterior Alpha Rhythm in Human EEG: Frequency Components, Their Cortical Sources, and Temporal Behavior. Scientific Reports, 7(1), 8249. 10.1038/s41598-017-08421-z

30. Park, J., Ho, R. L. M., Wang, W., Nguyen, V. Q., & Coombes, S. A. (2024). The effect of age on alpha rhythms in the human brain derived from source localized resting-state electroencephalography. NeuroImage, 292, 120614. 10.1016/j.neuroimage.2024.120614

31. Capilla, A., Schoffelen, J.-M., Paterson, G., Thut, G., & Gross, J. (2014). Dissociated α-Band Modulations in the Dorsal and Ventral Visual Pathways in Visuospatial Attention and Perception. Cerebral Cortex, 24(2), 550–561. 10.1093/cercor/bhs343

32. Sokoliuk, R., Mayhew, S. D., Aquino, K. M., Wilson, R., Brookes, M. J., Francis, S. T., Hanslmayr, S., & Mullinger, K. J. (2019). Two Spatially Distinct Posterior Alpha Sources Fulfill Different Functional Roles in Attention. The Journal of Neuroscience, 39(36), 7183– 7194. 10.1523/JNEUROSCI.1993-18.2019

33. Rodriguez-Larios, J., ElShafei, A., Wiehe, M., & Haegens, S. (2022). Visual Working Memory Recruits Two Functionally Distinct Alpha Rhythms in Posterior Cortex. Eneuro, 9(5), ENEURO.0159-22.2022. 10.1523/ENEURO.0159-22.2022

34. Cruz, G., Melcón, M., Sutandi, L., Matias Palva, J., Palva, S., & Thut, G. (2025). Oscillatory Brain Activity in the Canonical Alpha-Band Conceals Distinct Mechanisms in Attention. The Journal of Neuroscience, 45(1), e0918242024. 10.1523/JNEUROSCI.0918-24.2024

35. Zhou, Y. J., Van Es, M. W. J., & Haegens, S. (2025). Distinct alpha networks modulate different aspects of perceptual decision-making. PLOS Biology, 23(10), e3003461. 10.1371/journal.pbio.3003461

36. Tröndle, M., Popov, T., Pedroni, A., Pfeiffer, C., Barańczuk-Turska, Z., & Langer, N. (2021). *Decomposing age effects in EEG alpha power* [Preprint]. Neuroscience. 10.1101/2021.05.26.445765

37. Finley, A. J., Angus, D. J., Van Reekum, C. M., Davidson, R. J., & Schaefer, S. M. (2022). Periodic and aperiodic contributions to theta-beta ratios across adulthood. Psychophysiology, 59(11), e14113. 10.1111/psyp.14113

38. Turner, C., Baylan, S., Bracco, M., Cruz, G., Hanzal, S., Keime, M., Kuye, I., McNeill, D., Ng, Z., Van Der Plas, M., Ruzzoli, M., Thut, G., Trajkovic, J., Veniero, D., Wale, S. P., Whear, S., & Learmonth, G. (2023). Developmental changes in individual alpha frequency: Recording EEG data during public engagement events. Imaging Neuroscience, 1, 1–14. 10.1162/imag_a_00001

39. Van Dijk, H., Van Wingen, G., Denys, D., Olbrich, S., Van Ruth, R., & Arns, M. (2022). The two decades brainclinics research archive for insights in neurophysiology (TDBRAIN) database. Scientific Data, 9(1), 333. 10.1038/s41597-022-01409-z

40. Pritchard, W. S. (1992). The Brain in Fractal Time: 1/F-Like Power Spectrum Scaling of the Human Electroencephalogram. International Journal of Neuroscience, 66(1–2), 119–129. 10.3109/00207459208999796

41. Tenke, C., & Kayser, J. (2005). Reference-free quantification of EEG spectra: Combining current source density (CSD) and frequency principal components analysis (fPCA). Clinical Neurophysiology, 116(12), 2826–2846. 10.1016/j.clinph.2005.08.007

42. Barry, R. J., & De Blasio, F. M. (2017). EEG frequency PCA in EEG-ERP dynamics. Psychophysiology, 55(5), e13042. 10.1111/psyp.13042

43. Nakhnikian, A., Oribe, N., Hirano, S., Fujishima, Y., Hirano, Y., Nestor, P. G., Francis, G. A., Levin, M., & Spencer, K. M. (2024). Spectral decomposition of resting state electroencephalogram reveals unique theta/alpha activity in schizophrenia. European Journal of Neuroscience, 59(8), 1946–1960. 10.1111/ejn.16244

44. Mørup, M., & Hansen, L. K. (2012). Archetypal analysis for machine learning and data mining. *Neurocomputing*, Special Issue on Machine Learning for Signal Processing 2010, *80*, 54–63. 10.1016/j.neucom.2011.06.033

45. Cutler, A., & Breiman, L. (1994). Archetypal Analysis. Technometrics, 36(4), 338–347. 10.1080/00401706.1994.10485840

46. Walter, W. G. (1938). CRITICAL REVIEW: THE TECHNIQUE AND APPLICATION OF ELECTRO-ENCEPHALOGRAPHY. *Journal of Neurology*, Neurosurgery & Psychiatry, 1(4), 359–385. 10.1136/jnnp.1.4.359

47. Scally, B., Burke, M. R., Bunce, D., & Delvenne, J.-F. (2018). Resting-state EEG power and connectivity are associated with alpha peak frequency slowing in healthy aging. Neurobiology of Aging, 71, 149–155. 10.1016/j.neurobiolaging.2018.07.004

48. Cesnaite, E., Steinfath, P., Jamshidi Idaji, M., Stephani, T., Kumral, D., Haufe, S., Sander, C., Hensch, T., Hegerl, U., Riedel-Heller, S., Röhr, S., Schroeter, M. L., Witte, A., Villringer, A., & Nikulin, V. V. (2023). Alterations in rhythmic and non-rhythmic resting-state EEG activity and their link to cognition in older age. NeuroImage, 268, 119810. 10.1016/j.neuroimage.2022.119810

49. Getzmann, S., Gajewski, P. D., Schneider, D., & Wascher, E. (2024). Resting-state EEG data before and after cognitive activity across the adult lifespan and a 5-year follow-up. Scientific Data, 11(1), 988. 10.1038/s41597-024-03797-w

50. Babayan, A., Erbey, M., Kumral, D., Reinelt, J. D., Reiter, A. M. F., Röbbig, J., Schaare, H. L., Uhlig, M., Anwander, A., Bazin, P.-L., Horstmann, A., Lampe, L., Nikulin, V. V., Okon-Singer, H., Preusser, S., Pampel, A., Rohr, C. S., Sacher, J., Thöne-Otto, A., … Villringer, A. (2019). A mind-brain-body dataset of MRI, EEG, cognition, emotion, and peripheral physiology in young and old adults. Scientific Data, 6(1), 180308. 10.1038/sdata.2018.308

51. Clayton, M. S., Yeung, N., & Cohen Kadosh, R. (2018). The many characters of visual alpha oscillations. European Journal of Neuroscience, 48(7), 2498–2508. 10.1111/ejn.13747

52. Clark, C. R., Veltmeyer, M. D., Hamilton, R. J., Simms, E., Paul, R., Hermens, D., & Gordon, E. (2004). Spontaneous alpha peak frequency predicts working memory performance across the age span. International Journal of Psychophysiology, 9.

53. Grandy, T. H., Werkle-Bergner, M., Chicherio, C., Lövdén, M., Schmiedek, F., & Lindenberger, U. (2013). Individual alpha peak frequency is related to latent factors of general cognitive abilities. NeuroImage, 79, 10–18. 10.1016/j.neuroimage.2013.04.059

54. Klimesch, W., Schimke, H., & Schwaiger, J. (1994). Episodic and semantic memory: An analysis in the EEG theta and alpha band. Electroencephalography and Clinical Neurophysiology, 91(6), 428–441. 10.1016/0013-4694(94)90164-3

55. Corcoran, A. W., Alday, P. M., Schlesewsky, M., & Bornkessel-Schlesewsky, I. (2018). Toward a reliable, automated method of individual alpha frequency (IAF) quantification. Psychophysiology, 55(7), e13064. 10.1111/psyp.13064

56. Schaworonkow, N., & Nikulin, V. V. (2022). Is sensor space analysis good enough? Spatial patterns as a tool for assessing spatial mixing of EEG/MEG rhythms. NeuroImage, 253, 119093. 10.1016/j.neuroimage.2022.119093

57. Cohen, M. X. (2022). A tutorial on generalized eigendecomposition for denoising, contrast enhancement, and dimension reduction in multichannel electrophysiology. NeuroImage, 247, 118809. 10.1016/j.neuroimage.2021.118809

58. Pfurtscheller, G., Brunner, C., Schlögl, A., & Lopes Da Silva, F. H. (2006). Mu rhythm (de)synchronization and EEG single-trial classification of different motor imagery tasks. NeuroImage, 31(1), 153–159. 10.1016/j.neuroimage.2005.12.003

59. Wisniewski, M. G., Joyner, C. N., Zakrzewski, A. C., & Makeig, S. (2024). Finding tau rhythms in EEG: An independent component analysis approach. Human Brain Mapping, 45(2), e26572. 10.1002/hbm.26572

60. Cave, A. E., De Blasio, F. M., Chang, D. H., Münch, G. W., & Steiner-Lim, G. Z. (2025). Eyes-open and eyes-closed EEG of older adults with subjective cognitive impairment versus healthy controls: A frequency principal components analysis study. Brain Research, 1850, 149399. 10.1016/j.brainres.2024.149399

61. Barry, R. J., De Blasio, F. M., Duda, A. T., & Munford, B. S. (2025). Prestimulus EEG Oscillations and Pink Noise Affect Go/No-Go ERPs. Sensors, 25(6), 1733. 10.3390/s25061733

62. Knyazeva, M. G., Barzegaran, E., Vildavski, V. Y., & Demonet, J.-F. (2018). Aging of human alpha rhythm. Neurobiology of Aging, 69, 261–273. 10.1016/j.neurobiolaging.2018.05.018

63. Freschl, J., Azizi, L. A., Balboa, L., Kaldy, Z., & Blaser, E. (2022). The development of peak alpha frequency from infancy to adolescence and its role in visual temporal processing: A meta-analysis. Developmental Cognitive Neuroscience, 57, 101146. 10.1016/j.dcn.2022.101146

64. Menétrey, M. Q., Roinishvili, M., Chkonia, E., Herzog, M. H., & Pascucci, D. (2024). Alpha peak frequency affects visual performance beyond temporal resolution. Imaging Neuroscience, 2, 1–12. 10.1162/imag_a_00107

65. Ramsay, I. S., Lynn, P. A., Schermitzler, B., & Sponheim, S. R. (2021). Individual alpha peak frequency is slower in schizophrenia and related to deficits in visual perception and cognition. Scientific Reports, 11(1), 17852. 10.1038/s41598-021-97303-6

66. Catalano, L. T., Reavis, E. A., Wynn, J. K., & Green, M. F. (2024). Peak alpha frequency in schizophrenia, bipolar disorder, and healthy volunteers: Associations with visual information processing and cognition. Biological Psychiatry: Cognitive Neuroscience and Neuroimaging, 9(11), 1132–1140.

67. Murphy, M., & Öngür, D. (2019). Decreased peak alpha frequency and impaired visual evoked potentials in first episode psychosis. NeuroImage: Clinical, 22, 101693.

68. Tenke, C. E., Kayser, J., Alvarenga, J. E., Abraham, K. S., Warner, V., Talati, A., Weissman, M. M., & Bruder, G. E. (2018). Temporal stability of posterior EEG alpha over twelve years. Clinical Neurophysiology, 129(7), 1410–1417. 10.1016/j.clinph.2018.03.037

69. Arutiunian, V., Opdahl, M., Sullivan, C. A. W., Santhosh, M., Neuhaus, E., Borland, H., Bernier, R. A., Bookheimer, S. Y., Dapretto, M., Jack, A., Jeste, S., McPartland, J. C., Naples, A., Van Horn, J. D., Pelphrey, K. A., Webb, S. J., & Gupta, A. R. (2025). Number of Alpha Peaks in the Electroencephalogram Is Associated With Clinical Phenotype and Copy Number Variants in Youths With Autism. *Biological Psychiatry: Cognitive Neuroscience and Neuroimaging*, S2451902225003027. 10.1016/j.bpsc.2025.10.001

70. Chiang, A. K. I., Rennie, C. J., Robinson, P. A., Roberts, J. A., Rigozzi, M. K., Whitehouse, R. W., Hamilton, R. J., & Gordon, E. (2008). Automated characterization of multiple alpha peaks in multi-site electroencephalograms. Journal of Neuroscience Methods, 168(2), 396–411. 10.1016/j.jneumeth.2007.11.001

71. Ronconi, L., Busch, N. A., & Melcher, D. (2018). Alpha-band sensory entrainment alters the duration of temporal windows in visual perception. Scientific Reports, 8(1), 11810. 10.1038/s41598-018-29671-5

72. Lin, Y.-J., Shukla, L., Dugué, L., Valero-Cabré, A., & Carrasco, M. (2021). Transcranial magnetic stimulation entrains alpha oscillatory activity in occipital cortex. Scientific Reports, 11(1), 18562. 10.1038/s41598-021-96849-9

73. Trajkovic, J., Di Gregorio, F., Marcantoni, E., Thut, G., & Romei, V. (2022). A TMS/EEG protocol for the causal assessment of the functions of the oscillatory brain rhythms in perceptual and cognitive processes. STAR Protocols, 3(2), 101435. 10.1016/j.xpro.2022.101435

74. Millard, S. K., Speis, D. B., Skippen, P., Chiang, A. K. I., Chang, W., Lin, A. J., Furman, A. J., Mazaheri, A., Seminowicz, D. A., & Schabrun, S. M. (2024). Can non-invasive brain stimulation modulate peak alpha frequency in the human brain? A systematic review and meta-analysis. European Journal of Neuroscience, 60(3), 4182–4200. 10.1111/ejn.16424

75. D’Angelo, M., Lanfranco, R. C., Chancel, M., & Ehrsson, H. H. (2026). Parietal alpha frequency shapes own-body perception by modulating the temporal integration of bodily signals. Nature Communications, 17(1), 53. 10.1038/s41467-025-67657-w

76. Michel, C. M., & Bréchet, L. (2026). EEG microstates: From methodological foundations to clinical translation. Trends in Neurosciences, 49(6), 433–449. 10.1016/j.tins.2026.04.003

77. Capilla, A., Schoffelen, J.-M., Paterson, G., Thut, G., & Gross, J. (2014). Dissociated α-Band Modulations in the Dorsal and Ventral Visual Pathways in Visuospatial Attention and Perception. Cerebral Cortex, 24(2), 550–561. 10.1093/cercor/bhs343

78. Mahjoory, K., Cesnaite, E., Hohlefeld, F. U., Villringer, A., & Nikulin, V. V. (2019). Power and temporal dynamics of alpha oscillations at rest differentiate cognitive performance involving sustained and phasic cognitive control. NeuroImage, 188, 135–144. 10.1016/j.neuroimage.2018.12.001

79. Hülsdünker, T., & Mierau, A. (2021). Visual Perception and Visuomotor Reaction Speed Are Independent of the Individual Alpha Frequency. Frontiers in Neuroscience, 15, 620266. 10.3389/fnins.2021.620266

80. Buergers, S., & Noppeney, U. (2022). The role of alpha oscillations in temporal binding within and across the senses. Nature Human Behaviour, 6(5), 732–742. 10.1038/s41562-022-01294-x

81. Busch, N., Geyer, T., & Zinchenko, A. (2024). Individual peak alpha frequency does not index individual differences in inhibitory cognitive control. Psychophysiology, 61(8), e14586. 10.1111/psyp.14586

82. Cottier, T., Turner, W., Chae, V. J., Holcombe, A. O., & Hogendoorn, H. (2025). No Evidence That Resting-State Individual Alpha Frequency Represents a Mechanism Underlying Motion-Position Illusions. European Journal of Neuroscience, 62(9), e70250. 10.1111/ejn.70250

83. Arns, M. (2012). EEG-Based Personalized Medicine in ADHD: Individual Alpha Peak Frequency as an Endophenotype Associated with Nonresponse. Journal of Neurotherapy, 16(2), 123–141. 10.1080/10874208.2012.677664

84. Arns, M., Vollebregt, M. A., Palmer, D., Spooner, C., Gordon, E., Kohn, M., Clarke, S., Elliott, G. R., & Buitelaar, J. K. (2018). Electroencephalographic biomarkers as predictors of methylphenidate response in attention-deficit/hyperactivity disorder. European Neuropsychopharmacology, 28(8), 881–891. 10.1016/j.euroneuro.2018.06.002

85. Furman, A. J., Meeker, T. J., Rietschel, J. C., Yoo, S., Muthulingam, J., Prokhorenko, M., Keaser, M. L., Goodman, R. N., Mazaheri, A., & Seminowicz, D. A. (2018). Cerebral peak alpha frequency predicts individual differences in pain sensitivity. NeuroImage, 167, 203– 210. 10.1016/j.neuroimage.2017.11.042

86. Voetterl, H. T. S., Sack, A. T., Olbrich, S., Stuiver, S., Rouwhorst, R., Prentice, A., Pizzagalli, D. A., Van Der Vinne, N., Van Waarde, J. A., Brunovsky, M., Van Oostrom, I., Reitsma, B., Fekkes, J., Van Dijk, H., & Arns, M. (2023). Alpha peak frequency-based Brainmarker-I as a method to stratify to pharmacotherapy and brain stimulation treatments in depression. Nature Mental Health, 1(12), 1023–1032. 10.1038/s44220-023-00160-7

87. Wen, H., & Liu, Z. (2016). Separating Fractal and Oscillatory Components in the Power Spectrum of Neurophysiological Signal. Brain Topography, 29(1), 13–26. 10.1007/s10548-015-0448-0

88. Gao, R., Peterson, E. J., & Voytek, B. (2017). Inferring synaptic excitation/inhibition balance from field potentials. NeuroImage, 158, 70–78. 10.1016/j.neuroimage.2017.06.078

89. Donoghue, T., Haller, M., Peterson, E. J., Varma, P., Sebastian, P., Gao, R., Noto, T., Lara, A. H., Wallis, J. D., Knight, R. T., Shestyuk, A., & Voytek, B. (2020). Parameterizing neural power spectra into periodic and aperiodic components. Nature Neuroscience, 23(12), 1655– 1665. 10.1038/s41593-020-00744-x

90. Gerster, M., Waterstraat, G., Litvak, V., Lehnertz, K., Schnitzler, A., Florin, E., Curio, G., & Nikulin, V. (2022). Separating Neural Oscillations from Aperiodic 1/f Activity: Challenges and Recommendations. Neuroinformatics, 20(4), 991–1012. 10.1007/s12021-022-09581-8

91. Bai, S., Menétrey, M. Q., & Pascucci, D. (2026). What is next? Predictable visual sequences are encoded with anticipatory biases and reduced neural responses. iScience, 114697. 10.1016/j.isci.2026.114697

92. Delorme, A. (2023). EEG is better left alone. Scientific Reports, 13(1), 2372. 10.1038/s41598-023-27528-0

93. Delorme, A., & Makeig, S. (2004). EEGLAB: An open source toolbox for analysis of single-trial EEG dynamics including independent component analysis. Journal of Neuroscience Methods, 134(1), 9–21. 10.1016/j.jneumeth.2003.10.009

94. Mullen, T. R., Kothe, C. A. E., Chi, Y. M., Ojeda, A., Kerth, T., Makeig, S., Jung, T.-P., & Cauwenberghs, G. (2015). Real-time neuroimaging and cognitive monitoring using wearable dry EEG. IEEE Transactions on Biomedical Engineering, 62(11), 2553–2567. 10.1109/TBME.2015.2481482

95. Perrin, F., Pernier, J., Bertrand, O., & Echallier, J. F. (1989). Spherical splines for scalp potential and current density mapping. Electroencephalography and Clinical Neurophysiology, 72(2), 184–187. 10.1016/0013-4694(89)90180-6

96. Dien, J. (2010). The ERP PCA Toolkit: An open source program for advanced statistical analysis of event-related potential data. Journal of Neuroscience Methods, 187(1), 138–145. 10.1016/j.jneumeth.2009.12.009

97. Van De Ville, D., Farouj, Y., Preti, M. G., Liégeois, R., & Amico, E. (2021). When makes you unique: Temporality of the human brain fingerprint. Science advances, 7(42), eabj0751.

98. Finn, E. S., Shen, X., Scheinost, D., Rosenberg, M. D., Huang, J., Chun, M. M., Papademetris, X., & Constable, R. T. (2015). Functional connectome fingerprinting: Identifying individuals using patterns of brain connectivity. Nature Neuroscience, 18(11), Articolo 11. 10.1038/nn.4135

99. Tourbier, S., Rue Queralt, J., Glomb, K., Aleman-Gomez, Y., Mullier, E., Griffa, A., Schöttner, M., Wirsich, J., Tuncel, A., Jancovic, J., Bach Cuadra, M., & Hagmann, P. (2022). *connectomicslab/connectomemapper3: Connectome Mapper v3.1.0* (Versione v3.1.0) [Software]. Zenodo. 10.5281/ZENODO.7249263

100. Gramfort, A., Papadopoulo, T., Olivi, E., & Clerc, M. (2010). OpenMEEG: Opensource software for quasistatic bioelectromagnetics. BioMedical Engineering OnLine, 9(1), 45. 10.1186/1475-925X-9-45

101. Pagnotta, M. F., Pascucci, D., & Plomp, G. (s.d.). Selective attention involves a feature-specific sequential release from inhibitory gating. NeuroImage, 118782.

102. Pascucci, D., Hervais-Adelman, A., & Plomp, G. (2018). Gating by induced Α-Γ asynchrony in selective attention. Human Brain Mapping. 10.1002/hbm.24216

103. Grech, R., Cassar, T., Muscat, J., Camilleri, K. P., Fabri, S. G., Zervakis, M., Xanthopoulos, P., Sakkalis, V., & Vanrumste, B. (2008). Review on solving the inverse problem in EEG source analysis. Journal of neuroengineering and rehabilitation, 5(1), 1–33.

104. Daducci, A., Gerhard, S., Griffa, A., Lemkaddem, A., Cammoun, L., Gigandet, X., Meuli, R., Hagmann, P., & Thiran, J.-P. (2012). The connectome mapper: An open-source processing pipeline to map connectomes with MRI. PloS one, 7(12), e48121.

105. Desikan, R. S., Ségonne, F., Fischl, B., Quinn, B. T., Dickerson, B. C., Blacker, D., Buckner, R. L., Dale, A. M., Maguire, R. P., & Hyman, B. T. (2006). An automated labeling system for subdividing the human cerebral cortex on MRI scans into gyral based regions of interest. Neuroimage, 31(3), 968–980.

