## Supplementary Material for "The compositional nature of the human alpha rhythm"

#### Simulations

We assessed the performance of fPCA under three generative models representing distinct scenarios that can give rise to multiple spectral components: a single oscillatory process with inter-individual variability, between-participant heterogeneity, and within-participant mixtures of oscillatory processes. These models allowed us to test whether fPCA can distinguish individual-level mixtures of spectral components from patterns that may arise from variability across participants or from the mathematical constraints of the decomposition.

In the **single-source model (*single*)**, all participants expressed a single oscillatory source with a nominal peak frequency of 10 Hz, with individual peak frequencies varying around this value. This scenario represents a single underlying oscillatory process with inter-individual variability, as might occur for individual alpha peak frequency. Because PCA imposes orthogonality constraints, it can produce multiple components that do not correspond directly to the underlying generative factors, potentially introducing artificial structure even when the data are generated by a single latent process [1]. This model therefore provided a test of whether fPCA might identify multiple spectral components in the absence of multiple underlying oscillatory sources.

In the **between-participant mixture model (*between*)**, each participant expressed a single oscillatory source, but its nominal peak frequency was randomly assigned to one of three values: 9, 10, or 11 Hz. The sources were associated with different scalp distributions. This model represents population heterogeneity, in which distinct participant subgroups express different dominant peak frequencies without necessarily exhibiting mixtures of spectral components within individuals. It allowed us to test whether fPCA could produce multiple components from between-participant differences alone.

In the **within-participant mixture model (*within*)**, all three oscillatory sources, with nominal peak frequencies of 9, 10, and 11 Hz, were simultaneously expressed within each participant. Each source was assigned a distinct scalp distribution centered on electrodes Pz, PO8, and Cz (Figure S1). This model represents the coexistence of multiple oscillatory processes within individuals and corresponds to the type of generative scenario underlying our interpretation of multiple alpha-related spectral components. To assess the influence of spatial overlap on component recovery, we repeated this simulation with the three sources centered on nearby electrode locations (PO4, PO8, PO10).

For each model, we simulated EEG data from 200 participants, with 50 epochs of 2 s duration per participant, sampled at 250 Hz. Oscillatory sources were generated using autoregressive processes of order two (AR(2)), with coefficients determined by the source frequency and a damping parameter of ($r = 0.98$). Source signals were normalized before being projected onto the scalp using Gaussian spatial point-spread functions centered on the corresponding electrode locations. The spatial spread was controlled by ($\sigma = 100$ mm), with each spatial distribution normalized to its maximum. In the between- and within-participant mixture models, source frequencies were centered on their nominal values of 9, 10, and 11 Hz. To evaluate the effect of frequency variability, we varied the standard deviation of source peak frequencies across 11 conditions, from 0 to 2 Hz, including 10 logarithmically spaced nonzero values between 0.1 and 2 Hz. Frequency perturbations were sampled independently across participants and, in the within-participant model, across sources. The simulated oscillatory activity was combined with independently generated pink and white noise [2,3], each normalized to unit standard deviation.

The resulting synthetic EEG data were analyzed using the same fPCA pipeline applied to the empirical data (including the criterion of retaining components explaining at least 5% of total variance).

We evaluated the decomposition using three measures: the number of recovered components, the spatial dissimilarity between component score topographies within participants, and the relative expression in component topographic norms. Spatial dissimilarity was quantified as the mean of (1-|r|) across all pairs of component topographies, where (r) is their spatial correlation. This measure ranges from 0 for perfectly correlated or anticorrelated topographies to 1 for uncorrelated topographies, thus quantifying differences in topographic shape irrespective of polarity. Relative component expression was quantified for each participant as the difference between the maximum and mean component topographic norms, divided by the mean norm. This measure captures the extent to which the spatial expression of an individual's component topographies is dominated by a single component, with higher values indicating that the component with the largest norm stands out more strongly relative to the average across components. Both measures were calculated separately for each participant and summarized across participants using the mean and standard error of the mean for each frequency-variability condition.

Figure S1A shows the results of the main simulations. As the standard deviation of component peak frequencies across participants increased (Frequency SD), the number of estimated components increased in all three scenarios (first row). In the single-source model (blue), the number of components increased from one at low variability to four at high variability. In the between-participant mixture model (orange), it increased from three to four, while in the within-participant mixture model (yellow), it increased from two to five. These results show that increased variability in the underlying signals can lead fPCA to extract additional components, even when the generative process does not involve multiple oscillatory sources within individuals.

However, the three scenarios differed in the individual-level expression and spatial distribution of the recovered components. In the single-source and between-participant mixture models, the relative expression measure indicated a stronger dominance of one component, particularly as frequency variability increased (second row). By contrast, component expression remained more balanced in the within-participant mixture model. Thus, although fPCA extracted multiple components in all three scenarios, the individual-level expression of these components differed depending on the underlying generative process.

More importantly, the scenarios also differed in the spatial dissimilarity of the component score topographies (third row). In the single-source and between-participant mixture models, the recovered components showed relatively similar scalp topographies (low dissimilarity), whereas the within-participant mixture model yielded higher spatial dissimilarity, which remained comparatively high even at higher frequency variability. These findings suggest that inspecting component topographies, alongside their individual-level expression, is of key importance for distinguishing components that may arise from inter-individual variability from those reflecting multiple coexisting processes within individuals.

To further prove this point, Figure S1B compares the within-participant mixture model when the three generating sources had spatially distinct (blue) or highly overlapping (violet) scalp topographies. When the sources were spatially distinct, fPCA recovered multiple components even at relatively low levels of frequency variability. In contrast, when the source topographies overlapped strongly, the algorithm recovered only one component at low Frequency SD values, with the number increasing as frequency variability increased. However, this increase in the number of recovered components was not accompanied by a corresponding increase in spatial dissimilarity (third row). Missing values in the relative expression and spatial dissimilarity plots indicate conditions in which only one component was recovered, preventing these metrics from being computed. Together, these results highlight the importance of inspecting both the differences in component score topographies and their relative expression within individuals when interpreting fPCA decompositions of empirical data as well as validating with other methods, such as archetypal analysis and non-negative matrix factorization (see Supplementary Material: “Non-negative matrix factorization”).

#### Figure S1

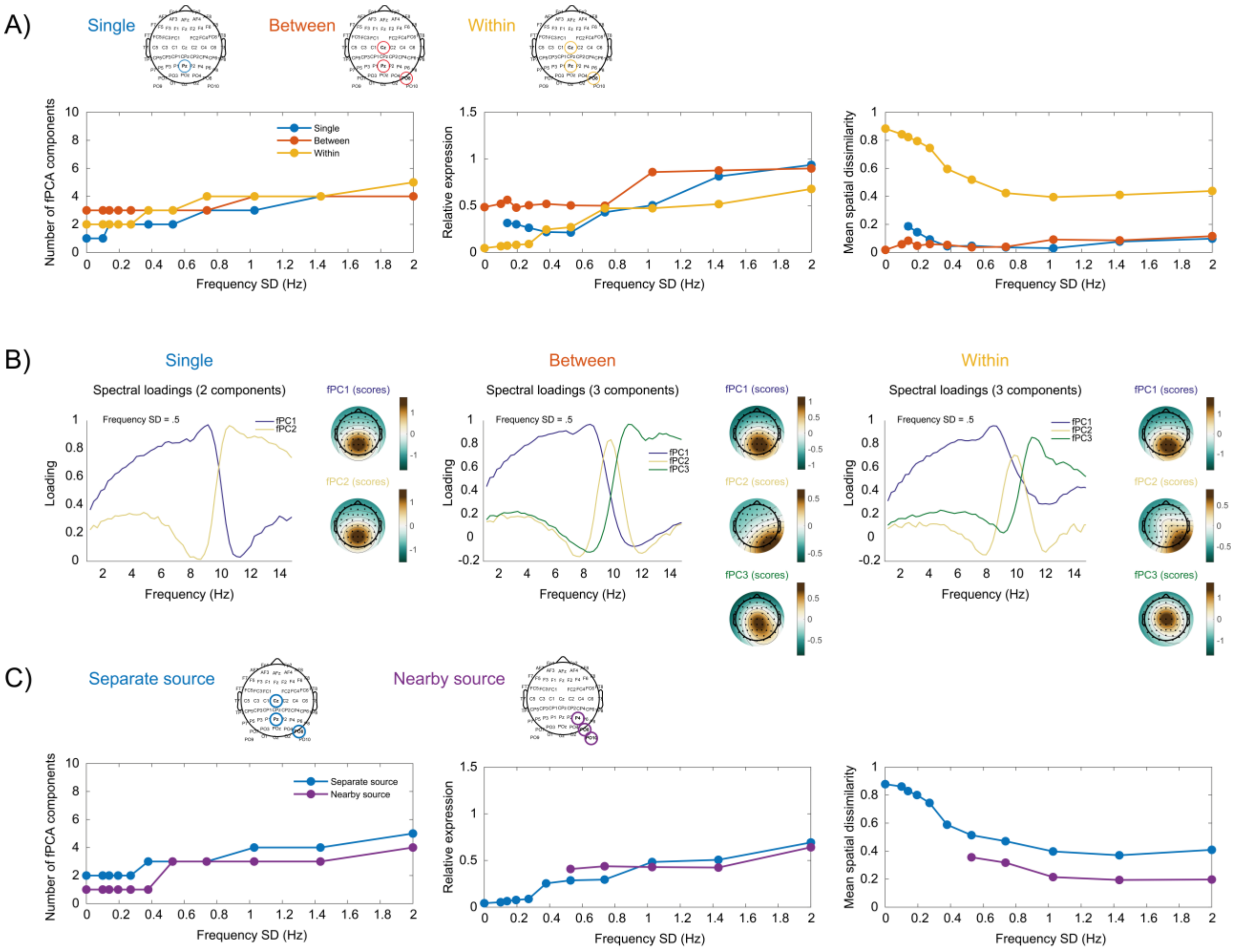


**Figure S1. Simulation of fPCA component recovery under different generative models and levels of frequency variability.** A) Single-source (blue), between-participant mixture (orange), and within-participant mixture (yellow) simulations. Topographies show the scalp distributions of the generating sources in each simulation. The left plot shows the number of recovered components as a function of the standard deviation of source peak frequencies across participants (Frequency SD), the middle plot shows the relative component expression metric as a function of Frequency SD, and the right plot shows the mean spatial dissimilarity. Dots indicate means across simulated participants. B) Estimated fPCA loadings and component score topographies for each generative model at an example Frequency SD of 0.5 Hz. C) Results for the within-participant mixture model with spatially distinct (blue) or highly overlapping (violet) generating source topographies. The topographies show the scalp locations of the generating sources for the spatially distinct and highly overlapping configurations.

#### Non-negative matrix factorization

As a complementary decomposition, we applied non-negative matrix factorization [NMF; 4] to the PSD data without z-scoring. Unlike fPCA, which decomposes mean-centered data into components with potentially positive and negative loadings, NMF constrains both the spectral components and their weights to be non-negative. This is consistent with the intrinsically non-negative nature of PSD and allows the observed spectra to be approximated as additive combinations of non-negative spectral components, without imposing orthogonality constraints.

NMF was applied to a matrix obtained by reshaping the data into subjects × electrodes by frequencies. Initialization was performed using non-negative double singular value decomposition [NNDSVD; 5] in Matlab. The number of components was determined from the cumulative proportion of squared singular values of the input matrix, retaining the smallest number of components accounting for at least 85% of the total squared singular-value energy. NMF was then fitted using a multiplicative-update algorithm, with a maximum of 1000 iterations and 100 replicates. The resulting factorization comprised spectral profiles and corresponding weights across subjects and electrodes. Scalp topographies were obtained by averaging component weights across subjects. NMF components were ordered based on the fPCA spectra to facilitate comparison. As shown in Figure S2, the NMF method consistently recovered four components with spectral profiles and topography differences well in line with the main results obtained with fPCA.

#### Figure S2

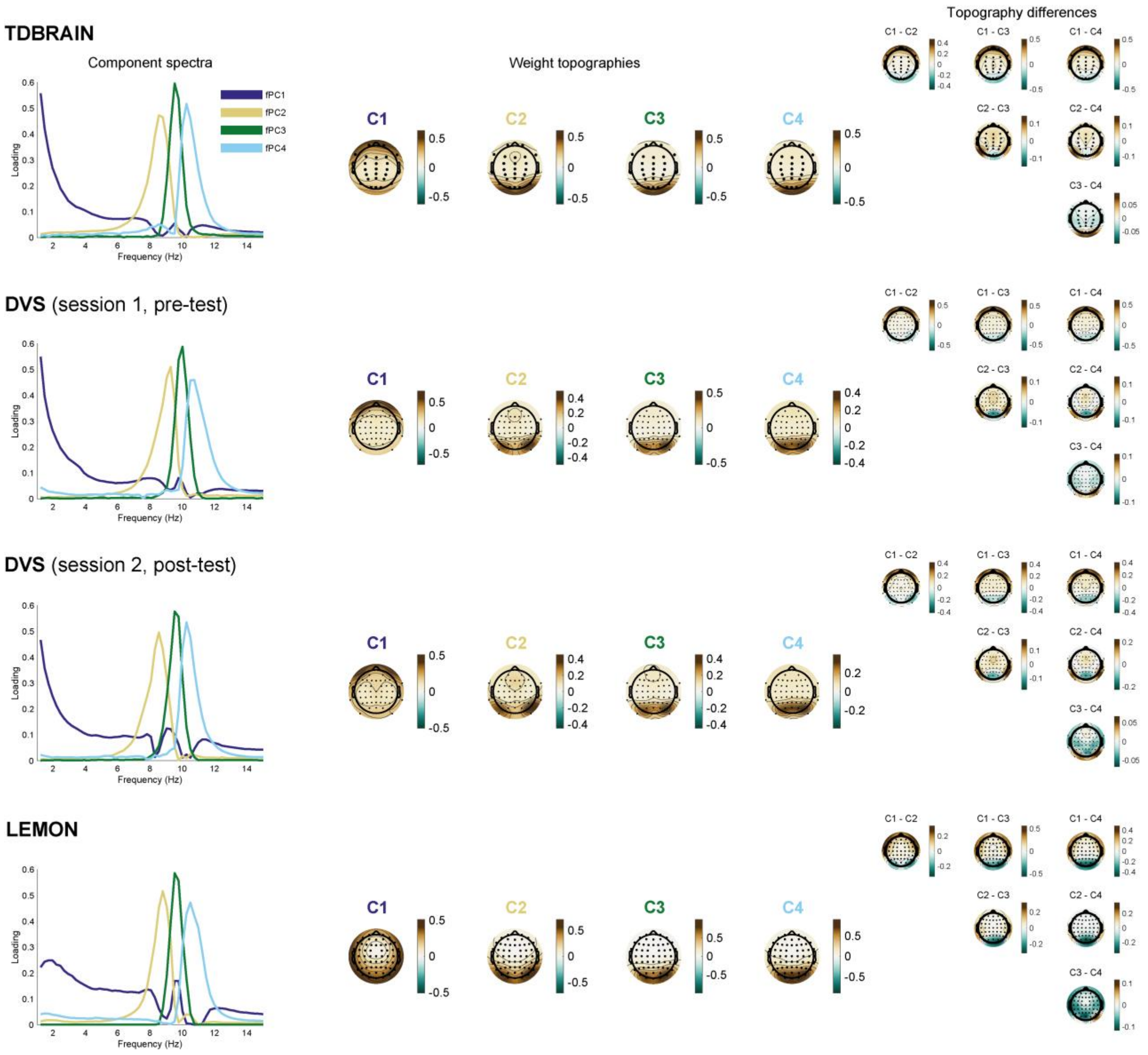


**Figure S2. Non-negative matrix factorization (NMF).** Results of NMF for four example datasets. For DVS, we report the first and last sessions, although all intermediate sessions produced similar results. The component spectra, topographic weights, and differences between topographies closely resemble the characteristics of the four main components identified in the main text using both fPCA and archetypal analysis. In particular, NMF highlights the distinction between slow and fast alpha components, which involve different temporal and parietal topographies.

#### Within-individual electrode subtyping

To further assess whether the alpha components observed in our fPCA reflect the coexistence of multiple spectral motifs within individuals, we performed an additional control analysis at the electrode level. In this analysis, we evaluated whether PSDs at individual electrodes were differentially associated with the three alpha components (slow, mid, and fast alpha). Prior to this analysis, we removed the contribution of the first fPCA component, which captured a broad low-frequency spectral pattern (Figures 1D, 2A, and 3A). For each subject and electrode, we regressed the PSD spectrum onto the loading profile of this component across frequencies and retained the residual spectrum. This step reduced the influence of slow, non-alpha spectral structure and allowed us to focus on the specific contribution of components in the alpha range to electrode-level PSD.

We then quantified the similarity between the residual PSD at each electrode and each of the three alpha components. For every subject, electrode, and component, we computed Pearson's correlation between the electrode PSD and the corresponding component loading profile across frequencies. Each electrode was assigned to the component with the highest positive correlation, provided that this correlation exceeded $r = 0.20$. Electrodes below this threshold were left unassigned. For each participant, we computed the proportion of electrodes assigned to each component, yielding a subject-by-component distribution of electrode assignments. These proportions provide a summary of the within-subject spatial expression of each alpha component. We further identified, for each participant, the two components with the highest electrode proportions and quantified the prevalence of these dominant component pairs across participants.

To visually verify that electrode assignments corresponded to the expected spectral patterns, we averaged the PSDs of electrodes assigned to each component within participants, z-scored the resulting spectra, and averaged them across participants sharing the same dominant component pair. As shown in Figure S3, electrodes assigned to different components exhibited spectral profiles consistent with the corresponding fPCA-derived alpha components. Furthermore, multiple alpha components were observed within the same participant, expressed by different subsets of electrodes and consistent with the distinct topographies associated with each component. The prevalence of dominant component pairs also varied across participants, with some combinations occurring more frequently than others.

### Figure S3

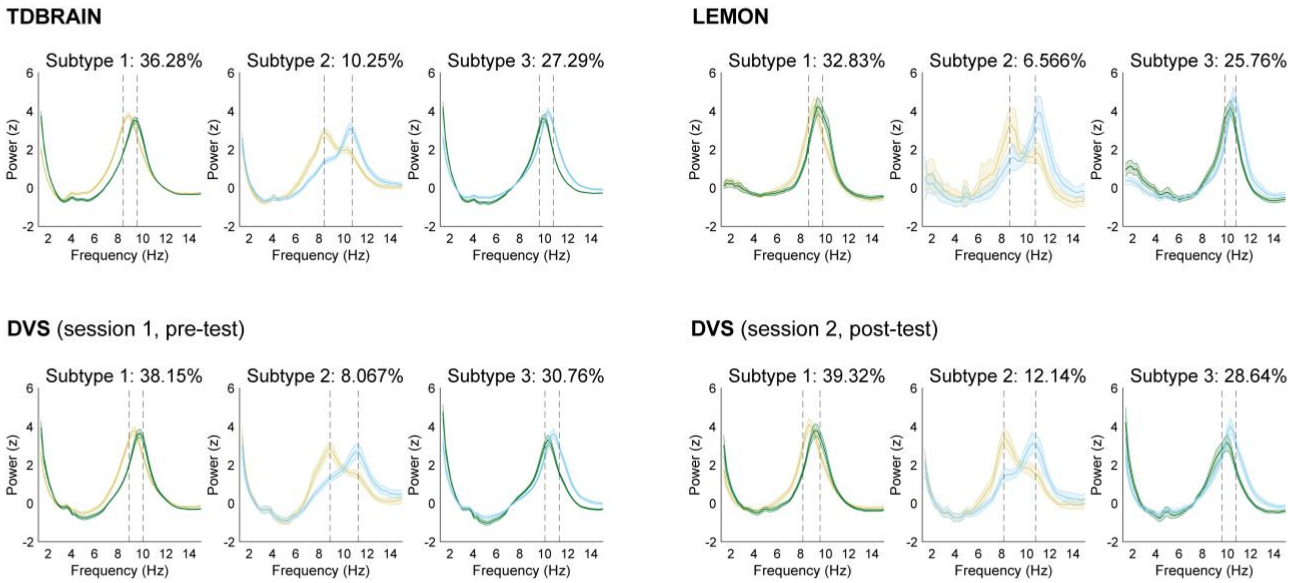


**Figure S3. Within-participant electrode subtyping across the main datasets.** Colors indicate the alpha components (slow, middle, and fast) using the same coding as in the main text. Dashed vertical lines indicate the peak frequencies of the two dominant fPCA components identified for each participant subgroup. The same procedure applied to the other sessions of the DVS dataset yielded highly comparable results.

### Figure S4

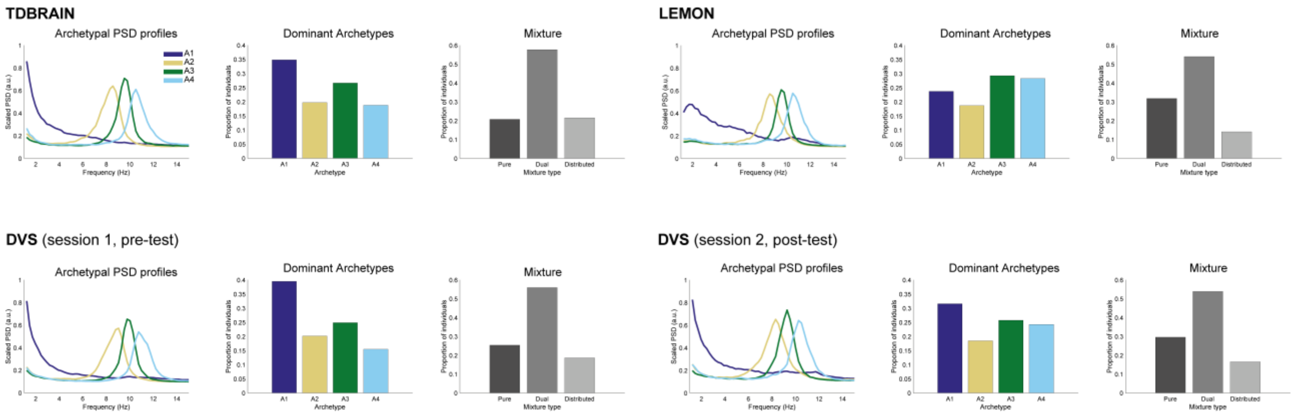


**Figure S4. Four archetypical components across datasets.** Results of archetypal analysis, including the recovered spectral archetypes, their dominance, and participation ratio metrics, as described in the main text, are shown for all datasets. For DVS, two example sessions are reported, as all other sessions yielded similar results.

### Figure S5

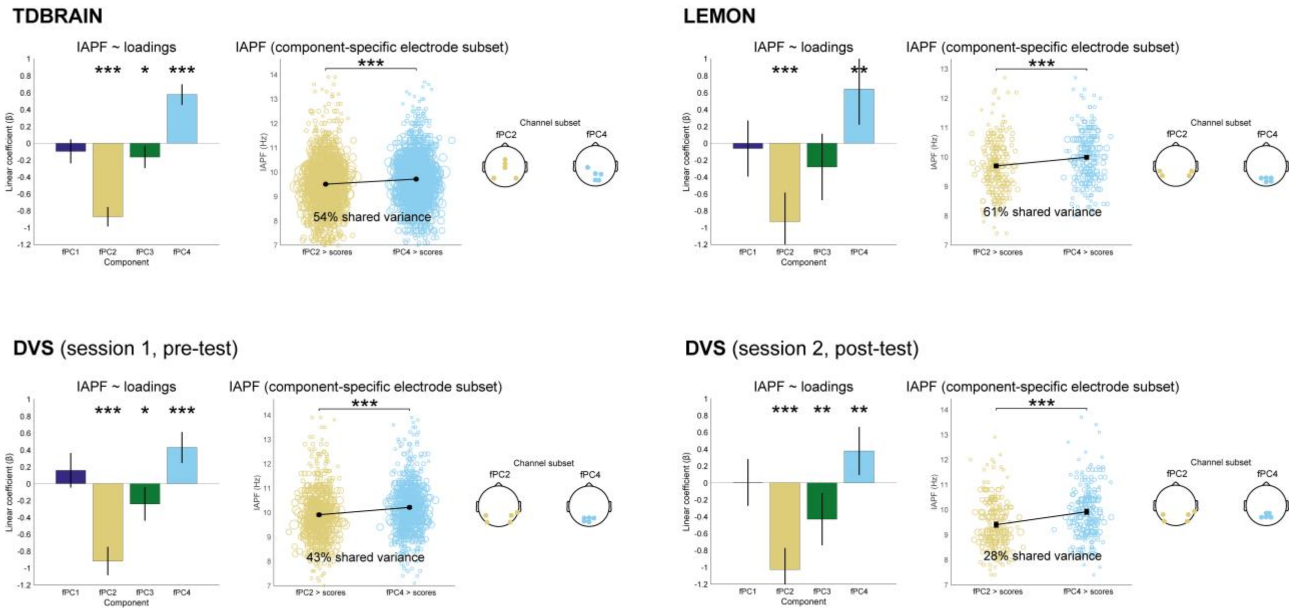


**Figure S5. Relation between alpha components and IAPF across datasets.** Linear models predicting IAPF from the scalp-averaged scores of the four fPCA spectral components, showing the consistent relationships between slow and fast alpha components and IAPF across all datasets, as reported in the main text. For DVS, two example sessions are shown, although the results were similar across the remaining sessions. The figure also shows IAPF at the electrode subsets expressing the slow (fPC2) and fast (fPC4) components most strongly, together with the electrode locations of the selected subsets for each dataset. Color coding is as in the main text. Significance levels: ***$p < .001$, **$p < .01$, *$p < .05$.

**Figure S6**

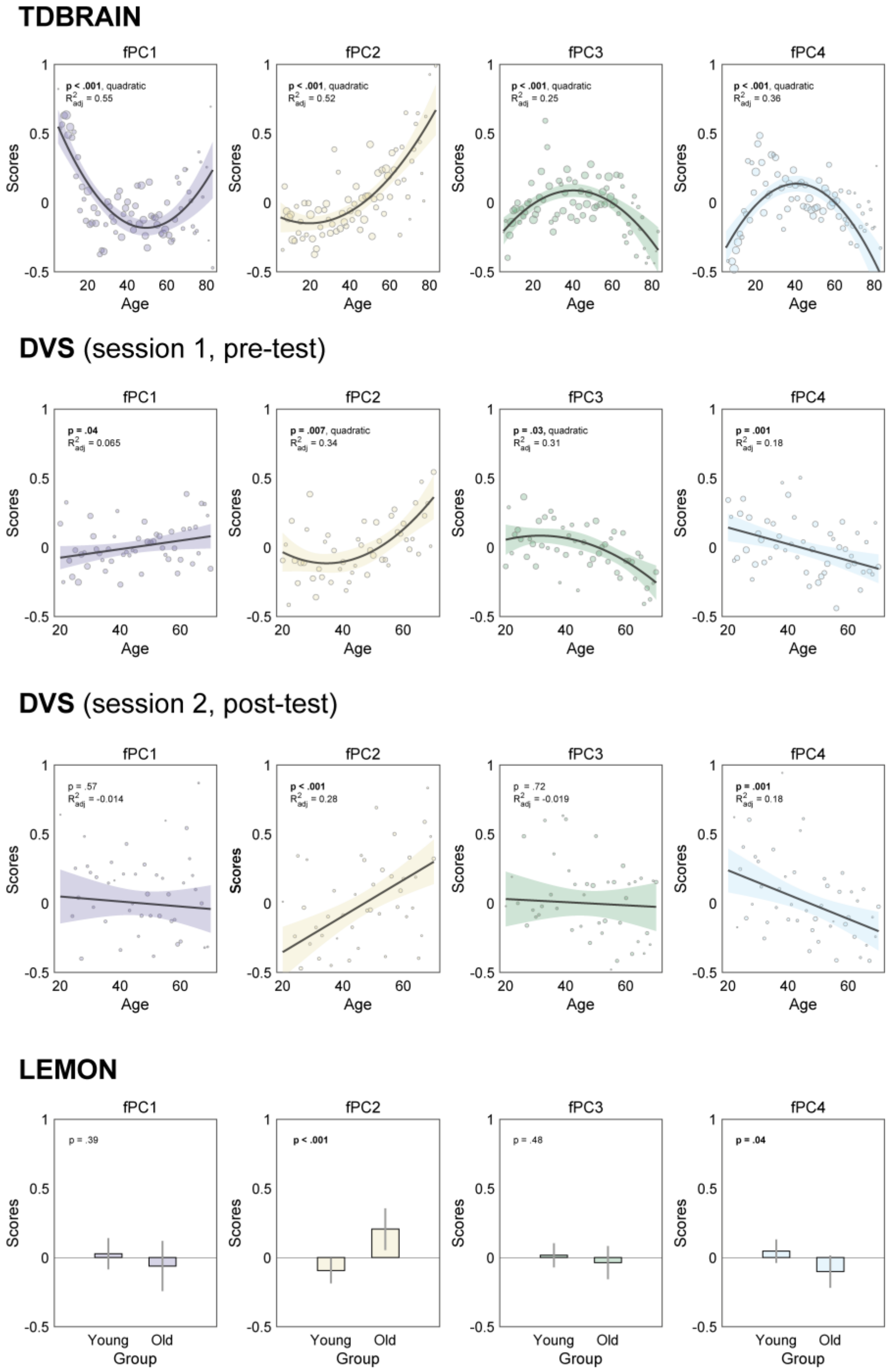


**Figure S6. Relation between alpha components and age across datasets.** Linear and quadratic models predicting age from the scalp-averaged scores of the four fPCA spectral components. Across datasets, the slow alpha component (fPC2) was among the components most strongly associated with age. Results for the other DVS sessions were similar. For LEMON, which included a group of younger (20–35 years) and older (59–77 years) participants, p-values are from independent-samples t-tests comparing the two age groups.

**Figure S7**

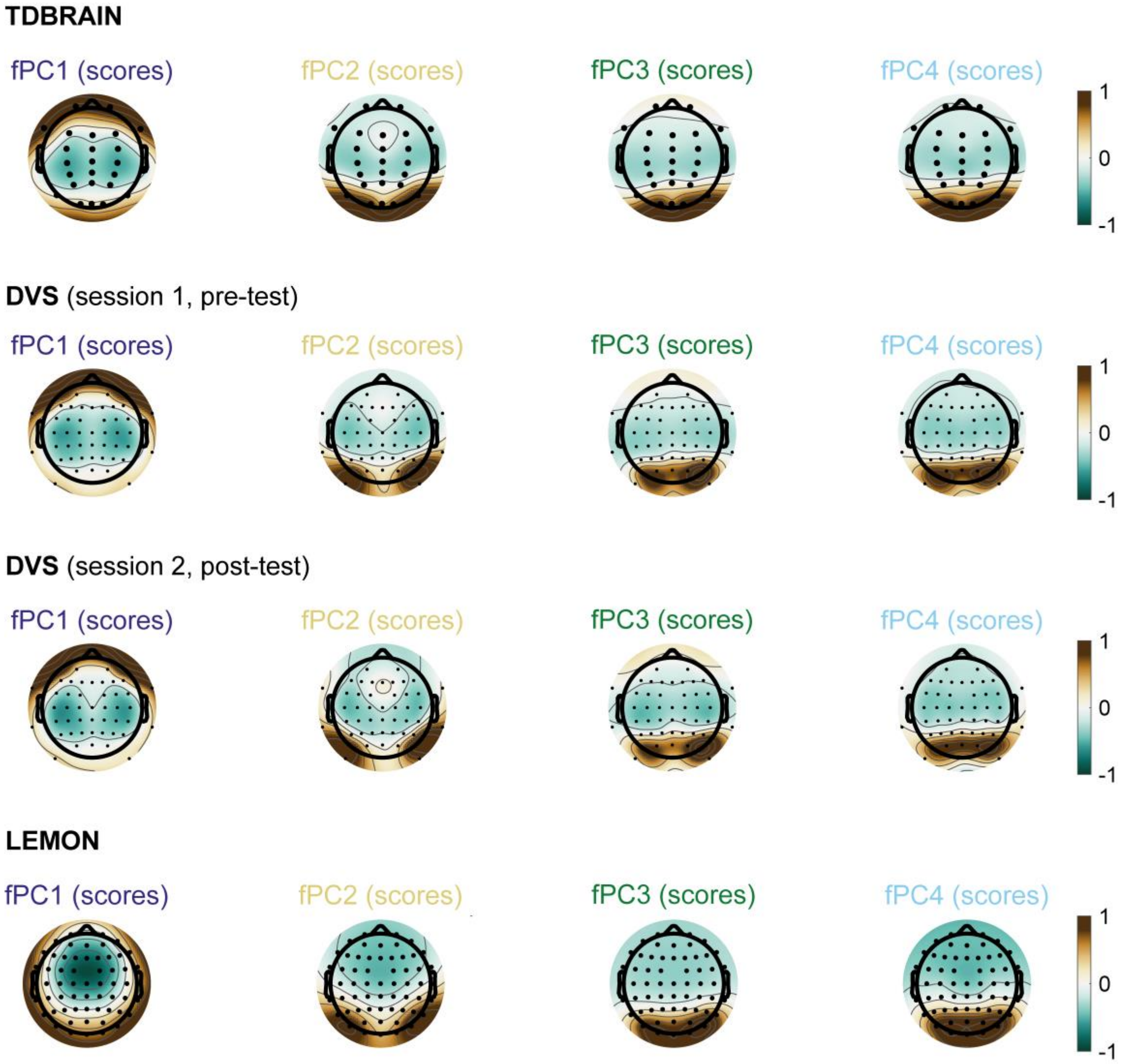


**Figure S7. Scalp topographies of component scores across datasets.** Scalp topographies associated with the four fPCA components are shown for the main datasets; similar topographies were observed across the other DVS sessions. The topographies highlight the distinction between fPC2 (slow alpha), with a stronger contribution from temporal electrodes, and fPC4 (fast alpha), with a stronger contribution from parieto-occipital electrodes. This distinction is particularly clear in datasets with denser electrode coverage (LEMON and DVS) but remains evident in TDBRAIN, despite its lower electrode density (see also Figure S2). LEMON topographies are reproduced from Figure 3C in the main text for comparison across datasets.

#### Replication of the four spectral components in a small sample dataset

To test whether the four components identified in large datasets can also be recovered in smaller, more typical EEG samples, we used an in-house dataset comprising RS-EEG recordings from 19 healthy participants aged 18 to 25 (8 females, 11 males), collected prior to participation in an independent study [6].

In this study, participants were recruited from École Polytechnique Fédérale de Lausanne (EPFL) and the University of Lausanne (UNIL) and received 30 CHF per hour for their participation. Two 2.5-minute resting-state EEG sessions were recorded—one with eyes open (EO) and one with eyes closed (EC)—using a 128-channel BioSemi ActiveTwo system (BioSemi, Amsterdam, the Netherlands). The cap was positioned to ensure that the A1 electrode was equidistant from the inion and nasion, as well as from both ears. EEG signals were sampled at 2048 Hz and referenced online to the common mode sense (CMS) and driven right leg (DRL) electrodes, with electrode voltages maintained within -20 μV to 20 μV. Offline preprocessing of the EC recordings included downsampling to 250 Hz, band-pass filtering between 0.5 and 40 Hz, and segmentation into 4-second epochs. The automatic cleaning to remove bad channels, epochs, and independent components identified via ICA, as well as the final preprocessing steps (electrode interpolation and average re-referencing) were similar to those used in the TDBRAIN and DVS datasets. Across the 2.5 minutes of EC recordings, an average of 1.6 electrodes, 0.6 trials, and 0.7 independent components were discarded.

Despite the much smaller number of participants, four spectral components, closely resembling those reported across the large datasets, were recovered from EC recordings using both fPCA (88.4% of variance explained; Figure S8) and archetypal analysis (95.4% of variance explained). This sample dataset used to illustrate the application of the code, together with the analysis code and derived results, is available in an online repository (DOI: https://doi.org/10.5281/zenodo.23600677).

### Figure S8

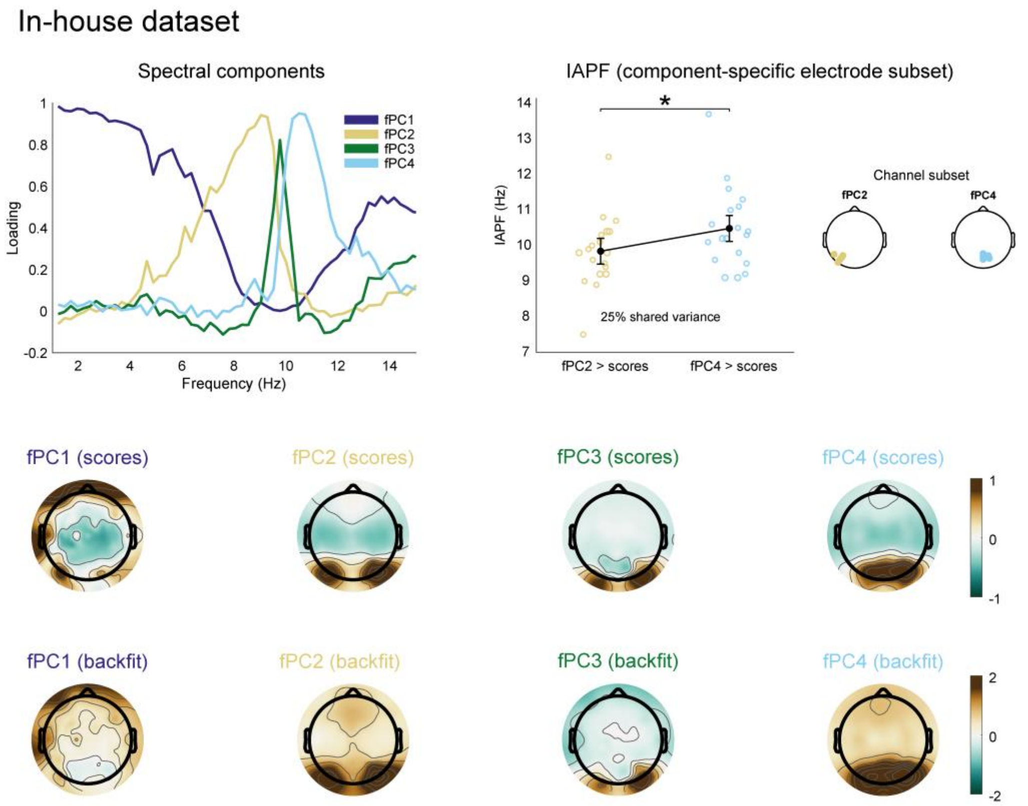


**Figure S8. Spectral decomposition of a small-sample dataset.** fPCA applied to an in-house dataset of 19 participants. Spectral loadings, scalp topographies of component scores, and IAPF estimated from electrode subsets expressing fPC2 (slow alpha) and fPC4 (fast alpha) most strongly are shown. The analyses followed the same approach as in the main text and revealed spectral components and topographies consistent with the main findings.

#### Spectral decomposition of eyes-open RS recording

The four components systematically identified using the fPCA approach and archetypal analysis across all datasets (e.g., Figures 1D-E) were consistently derived from eyes-closed (EC) recordings. Here, we applied the same pipeline to eyes-open (EO) recordings to assess whether these distinct spectral components could be replicated under different resting-state conditions.

We first preprocessed the EO recordings from the TDBRAIN dataset using a similar procedure to that applied to the EC recordings (see Methods). EO recordings were available for 1295 participants. On average, 2.19 electrodes, 23% of the data, and 0.9 independent components were removed during preprocessing. In addition, 72 participants were excluded due to insufficient recording duration and 43 participants for whom IAPF detection using FOOOF was unsuccessful. This resulted in a final sample of 1180 retained participants.

We next preprocessed the EO recordings from the DVS dataset, considering the 3-min segments preceding the cognitive task in the first session (608 participants) and, when available, the corresponding 3-min segments from the second session, recorded approximately 5 years later (208 participants). The recordings were preprocessed using a procedure similar to that applied to the EC recordings (see Methods). An average of 4.1 electrodes, 33.5% of the data, and 6.5 independent components were discarded across the 6 min of EO recordings. Moreover, participants were excluded if the automated pipeline retained less than 1 min of usable data in either session (29 participants in Session 1 and 15 in Session 2) or if no IAPF could be detected using FOOOF (37 participants in Session 1 and 12 in Session 2). This resulted in 542 participants retained for Session 1 and 181 participants retained for Session 2.

Finally, for the LEMON dataset, which had already been preprocessed (see Methods), we only performed interpolation of removed electrodes, re-referencing to the average reference, and segmentation into 4-second epochs. Of the 200 participants with available EO recordings, 185 were retained after excluding 15 participants for whom no IAPF could be identified.

Similarly to the fPCA results obtained from EC recordings, we consistently identified four components in the EO recordings from the TDBRAIN, DVS (both sessions separated by five years) and LEMON datasets (Figure S9). In all cases, these components jointly explained between 78% and 82% of the variance, with an overall higher contribution from the non-alpha component. The components in the alpha-band differed from that observed in EC recordings: only two components were found, corresponding to the slow and fast alpha components. The fourth component peaked at higher frequencies (approximately 20–30 Hz), in a range more consistent with beta oscillations. Regarding their scalp topographies, the slow and fast alpha components were relatively similar to those identified in EC recordings (e.g., Figure 3C), with the slow alpha component showing a more temporal distribution and the fast alpha component exhibiting a more occipitoparietal distribution.

The presence of two main alpha components is consistent with recent work using EO resting-state EEG recordings [7,8]. One possible explanation is that the middle alpha component, which showed the most pronounced occipital distribution in EC recordings (e.g., Figure 3E), may be less distinguishable in EO recordings because of its direct association with visual processing.

### Figure S9

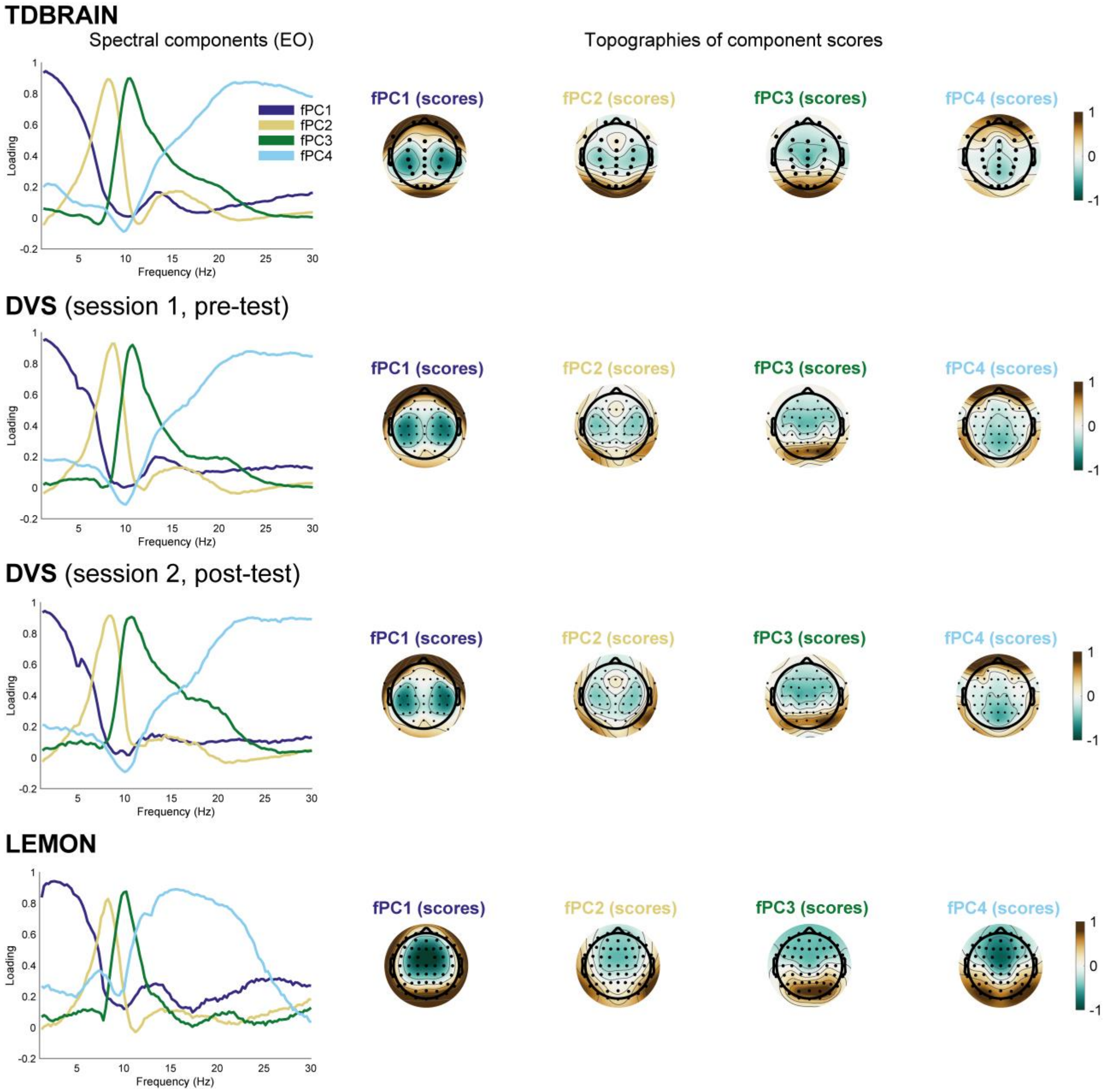


**Figure S9. fPCA of eyes-open (EO) resting-state EEG.** Spectral loadings and scalp topographies of component scores obtained from EO resting-state EEG recordings across datasets. Results for the other DVS sessions were similar. The analysis consistently identified a non-alpha component, slow and middle alpha components, and a higher-frequency, beta-like component, while the fast alpha component was absent across datasets.

#### Reference

1. Shinn, M. (2023). Phantom oscillations in principal component analysis. *Proceedings of the National Academy of Sciences*, *120*(48), e2311420120. https://doi.org/10.1073/pnas.2311420120
2. Barzegaran, E., Bosse, S., Kohler, P. J., & Norcia, A. M. (2019). EEGSourceSim: A framework for realistic simulation of EEG scalp data using MRI-based forward models and biologically plausible signals and noise. *Journal of Neuroscience Methods*, *328*, 108377. https://doi.org/10.1016/j.jneumeth.2019.108377
3. Haufe, S., & Ewald, A. (2019). A Simulation Framework for Benchmarking EEG-Based Brain Connectivity Estimation Methodologies. *Brain Topography*, *32*(4), 625–642. https://doi.org/10.1007/s10548-016-0498-y
4. Lee, D. D., & Seung, H. S. (1999). Learning the parts of objects by non-negative matrix factorization. *Nature*, *401*(6755), 788–791. https://doi.org/10.1038/44565
5. Boutsidis, C., & Gallopoulos, E. (2008). SVD based initialization: A head start for nonnegative matrix factorization. *Pattern Recognition*, *41*(4), 1350–1362. https://doi.org/10.1016/j.patcog.2007.09.010
6. Bai, S., Menétrey, M. Q., & Pascucci, D. (2026). What is next? Predictable visual sequences are encoded with anticipatory biases and reduced neural responses. *iScience*, 114697. https://doi.org/10.1016/j.isci.2026.114697
7. Barry, R. J., De Blasio, F. M., Duda, A. T., & Munford, B. S. (2025). Prestimulus EEG Oscillations and Pink Noise Affect Go/No-Go ERPs. *Sensors*, *25*(6), 1733. https://doi.org/10.3390/s25061733
8. Cave, A. E., De Blasio, F. M., Chang, D. H., Münch, G. W., & Steiner-Lim, G. Z. (2025). Eyes-open and eyes-closed EEG of older adults with subjective cognitive impairment versus healthy controls: A frequency principal components analysis study. *Brain Research*, *1850*, 149399. https://doi.org/10.1016/j.brainres.2024.149399
